# Haplotype-phased genomes of the barley leaf rust pathogen reveal evidence of repeat element expansion and somatic hybridization

**DOI:** 10.64898/2026.01.30.702850

**Authors:** Rebecca Spanner, Eric S. Nazareno, Eva C. Henningsen, Alexis Feist, Matthew J. Moscou, Matthew N. Rouse, Hayley Mangelson, Kyle Langford, Ivan Liachko, Jibril Lubega, Kostya Kanyuka, Jana Sperschneider, Peter N. Dodds, Melania Figueroa, Brian J. Steffenson

## Abstract

Barley leaf rust disease, caused by *Puccinia hordei*, leads to substantial yield losses and diminished malting quality of barley across temperate growing regions worldwide. To address the paucity of high-resolution genomic resources for this pathogen, we generated haplotype-phased, chromosome-scale assemblies for ten globally distributed isolates using PacBio HiFi and Hi-C sequencing. Phylogenomic analysis revealed seven distinct lineages of *P. hordei*, including evidence of nuclear exchange, with a shared nuclear haplotype detected between two US lineages. Nuclear genome sizes ranged from ∼140-147Mbp, with the exception of isolate 90ISR03 from Israel (∼163Mbp), which also harbored a 6.2Mbp supernumerary scaffold in one nucleus exhibiting chromosomal characteristics. Consistent with its larger genome, *P. hordei* had a higher repeat content (∼70%) than related cereal rust fungi, driven primarily by the proliferation of LTR retroelements and DNA transposons. Across the global pan-genome of 13 unique nuclear haplotypes, approximately one-third of all protein orthogroups were conserved across all isolates. Only 18% of predicted effector orthogroups were shared between all haplotypes, reflecting the highly dynamic and variable nature of the effector repertoire. The long-term propagation of clonal *P. hordei* lineages is apparent both within the US and globally, and nuclear exchange is important for generating novel diversity and virulence profiles. The high degree of genome plasticity is evident in extensive structural variation, including large-scale translocations and inversions as well as a putative accessory chromosome. These chromosome-level, haplotype-resolved genomes provide a foundational resource for exploring the evolution, diversity, and avirulence gene repertoire of *P. hordei*.

## Introduction

Barley leaf rust, caused by *Puccinia hordei* Otth, is the most important rust disease affecting barley (*Hordeum vulgare* L.) worldwide and is prevalent across all major barley-growing regions (Clifford, 1985; Park et al., 2015). Under conditions of high inoculum pressure, susceptible cultivars can suffer yield losses of up to 60% (Park et al., 2015). As an obligate biotroph, *P. hordei* has evolved to infect both wild (*Hordeum vulgare* subsp. *spontaneum*) and cultivated barley (*Hordeum vulgare* subsp. *vulgare*), subverting host immunity and acquiring nutrients via specialized feeding structures called haustoria (Anikster & Wahl, 1979; Parlevliet & Kievit, 1986).

While the disease is best managed by the deployment of resistant cultivars, the pathogen can overcome host resistance through virulence evolution driven by somatic hybridization, meiotic recombination, and spontaneous mutation (Anikster, 1984a; Figueroa et al., 2020). This has led to “boom and bust” cycles of deployed resistance genes in barley cultivars. For instance, virulence for a widely effective barley leaf rust gene in the United States, *Rph7.g*, has been observed since the 1990s due to its large-scale deployment in the eastern part of the country (Steffenson, 1993). Virulence for other key *Rph* genes has also increased based on 30-year survey data, highlighting the evolutionary potential of this pathogen (Nazareno et al., 2023).

*P. hordei* is believed to have coevolved in the Near East region of the Fertile Crescent to infect species of the primary host *Hordeum* and the alternate hosts of *Ornithogalum*, *Leopoldia*, and *Dipcadi*, which are members of the sub-family Scilloideae in the family Asparagaceae (d’Oliveira, 1960). Sexual reproduction of *P. hordei* on the alternate hosts has largely been limited to parts of the Middle East (such as Israel) (Anikster, 1984a), southern Europe (Anikster, 1982), and northern Africa (Anikster, 1982, 1984a) where both primary and alternate hosts are present in combination with the optimal conditions for teliospore germination (Anikster & Wahl, 1979; Savile, 1984; Yahyaoui, 1987). In North America, *Ornithogalum* is present in the eastern part of the continent; however, the aecial stage has not been recorded on this alternate host (Savile, 1984). Distinct pathotypes of *P. hordei* have been observed for uredinial isolates that infect both *H. vulgare* subsp. *vulgare* and *H. vulgare* subsp. *spontaneum* but this is not the case for uredinial isolates that infect *H. bulbosum* and *H. marinum* (Anikster, 1989). For most of its life cycle, *P. hordei* survives as dikaryotic spores, either asexual repeating urediniospores, overwintering teliospores or sexually-derived aeciospores from *Ornithogalum* (Anikster, 1989). The dikaryotic stage, characterized by the coexistence of two genetically distinct haploid nuclei per cell, is a unique feature of fungi, particularly basidiomycetes (Kruzel & Hull, 2010).

However, dikaryons pose a challenge for *de novo* genome assembly in basidiomycete fungi since conventional graph-based assemblers are unable to distinguish the origins of reads from large genomic regions of high similarity between homologous chromosomes from distinct nuclei (Zhang et al., 2022). This has previously resulted in genome assemblies with high proportions of collapsed regions. For rust fungi with large genomes and high repetitive content, approaches employing long reads alone often failed to completely phase the two nuclear genomes present in the dikaryon (Chen et al., 2019; Miller et al., 2018; Schwessinger et al., 2018). A milestone for *Puccinia* genome assembly has been the combined use of long reads and chromatin contact sequencing to fully phase the two nuclear genomes in the dikaryon (Li et al., 2019) with increased accuracy of PacBio high fidelity (HiFi) reads facilitating accurate phased assemblies (Duan et al., 2022). Information regarding the 3D structure of chromatin (HiC) within its constituent nucleus and where it contacts facilitates both nuclear phasing and scaffolding of individual chromosomes (Burton et al., 2013; Marie-Nelly et al., 2014). The genome assembler hifiasm uses PacBio HiFi reads to produce a phased assembly graph and can integrate HiC data to improve phasing (Cheng et al., 2021). With the aid of HiC-scaffolders (Ghurye et al., 2017) or other scaffolders using chromatin contact sequencing data, fully phased telomere-to-telomere nuclear assemblies are now possible. Haplotype-resolved genome assemblies have now been achieved in multiple dikaryotic rust species including *Austropuccinia psidii* (Luo et al., 2025), *Puccinia hordei* (Yu et al., 2025), *Puccinia graminis* f. sp. *tritici* (Li et al., 2019)*, Puccinia triticina* (Duan et al., 2022; Sperschneider et al., 2023)*, Puccinia coronata* f. sp. *avenae* (Henningsen et al., 2022, 2024, 2025)*, Puccinia striiformis* f. sp. *tritici* (Holden et al., 2025; Tam et al., 2025; Wang et al., 2024), and *Puccinia polysora* f. sp. *zeae* (Liang et al., 2023).

The development of haplotype-phased genome assemblies has revealed evidence of historical nuclear exchange events in rust fungi. The Ug99 wheat stem rust lineage emerged via somatic hybridization, with one haploid nuclear genome (hap01) derived from the Pgt21 lineage and the other (hap03) from a member of the Clade II lineage (Guo et al., 2022; Li et al., 2019; Spanner et al., 2025). With phased genomes, the movement of entire nuclei can be tracked (“haplotracking”) to evaluate the contribution of somatic hybridization to global rust population dynamics, particularly where sexual recombination is rare (Henningsen et al., 2024; Sperschneider et al., 2023). Nuclear exchange events have also been detected in *P. triticina* (Sperschneider et al., 2023), *P. coronata* f. sp. *avenae* (Henningsen et al., 2024, 2025) and *P. striiformis* f. sp. *tritici* (Holden et al., 2025), where they have also led to the emergence of strains with distinct virulence profiles that have become epidemiologically dominant. The detection of shared nuclei on distinct continents also implied long-distance dispersal of haplotypes through migration of members of clonal lineages and subsequent somatic hybridization.

The transfer of entire nuclei is an effective way for rust pathogens to obtain new virulence profiles through novel combinations of avirulence gene variants. In rust fungi, cloned avirulence (Avr) genes encode small proteinaceous effectors that are secreted during invasion of their host from specialized feeding structures called haustoria (Henningsen et al., 2025). Rust effectors facilitate infection by manipulating host targets, suppressing the host immune system, and aiding in nutrient absorption (Lorrain et al., 2019). When these proteins are recognized by plant receptors (directly or indirectly), an immune response is triggered leading to cell death (Dodds, 2023; Dodds & Rathjen, 2010). This prevents these biotrophic pathogens from spreading through the plant and is the basis of most forms of genetic resistance used for disease management. The complete phasing of nuclear genomes in dikaryotic rust fungi will aid in the resolution of putative effector genotypes (Figueroa et al., 2023), as demonstrated in the development of the wheat stem rust Avr gene atlas (Spanner et al., 2025). It also enables resolution of loci residing in repeat-rich regions such as the mating type loci, which may influence sexual or asexual compatibility (Henningsen et al., 2024; Sperschneider et al., 2023). Such resources further support the generation of effector libraries that can be used for high-throughput screens to identify new Avr effector genes (Arndell et al., 2024; Chen et al., 2025; Spanner et al., 2025).

The lack of high-quality genome resources has limited evolutionary studies in *P. hordei* and has hampered the identification of Avr genes. In the present study, we sought to develop annotated haplotype-phased genomes of ten global isolates of *P. hordei* in order to investigate genome architecture and explore the relationships among these global *Ph* lineages.

## Materials and Methods

### Rust isolate preparation and phenotyping

Ten isolates of *P*. *hordei* were selected for sequencing in this study. Six isolates were from the U.S. states of Minnesota (17MN32B), Virginia (82VA01 and 93VA27), Texas (94TX04), Washington (92WA74), and California (15CA06C) and four isolates were from the countries of China (97CHN01), Germany (91DEU05), the Netherlands (91NLD202) and Israel (90ISR03) (**Figure 1A, Table S1**). The criteria used for selection of these isolates included their geographic origin, virulence phenotypes on the barley leaf rust differential set (Martin et al., 2020), and previous genotyping with AFLP markers (Sun et al., 2007). The domestic and foreign isolates were initially increased on the susceptible barley cultivar ‘Robust’ (PI 476976) in the greenhouse and in the Biosafety Level-3 (BSL-3) containment facility, respectively. Single uredinia were isolated, then increased to harvest spores for DNA extraction and phenotyping on the differential barley lines (Martin et al., 2020). Increased urediniospores were desiccated and then stored at - 80°C until needed. For the virulence phenotyping assays, isolates were heat shocked at 45°C for 15 min, re-humidified in a 80% relative humidity chamber for several hours, suspended in a lightweight mineral oil (Soltrol 170, Chevron Phillips Chemical Company, The Woodlands, TX), and then applied to seven to eight-day old plants at a rate of 5 mg per 500 µl of oil per tray (15 lines with 3-5 plants per line). After volatilization of the oil carrier, inoculated plants were placed in mist chambers overnight (∼16 hrs) to provide optimal conditions for infection by *P. hordei* (Martin et al., 2020). After the infection period, plants were allowed to air dry and moved to a greenhouse or growth chamber within the BSL-3 (foreign isolates) with day/night temperatures of 18/21°C and 16/8 hrs of light/dark cycle. Rust virulence phenotypes on the barley differentials were made at 8-10 days post inoculation (dpi) based on a 0-4 scale (Levine & Cherewick, 1952), where ratings of 0-2 were considered indicative of pathogen avirulence, and 3-4 of virulence. Virulence phenotypes were determined on the Bowman introgression lines carrying genes *Rph1-Rph15* (**Table S3**) (Martin et al., 2020). Isolates were also phenotyped for virulence on the original sources of the leaf resistance genes *Rph1* to *Rph24* (**Table S2**). Phenotypic scores based on the 0-4 scale were converted to a 0-9 scale to achieve a finer level of resolution for pathogen virulence levels.

**Figure 1.**
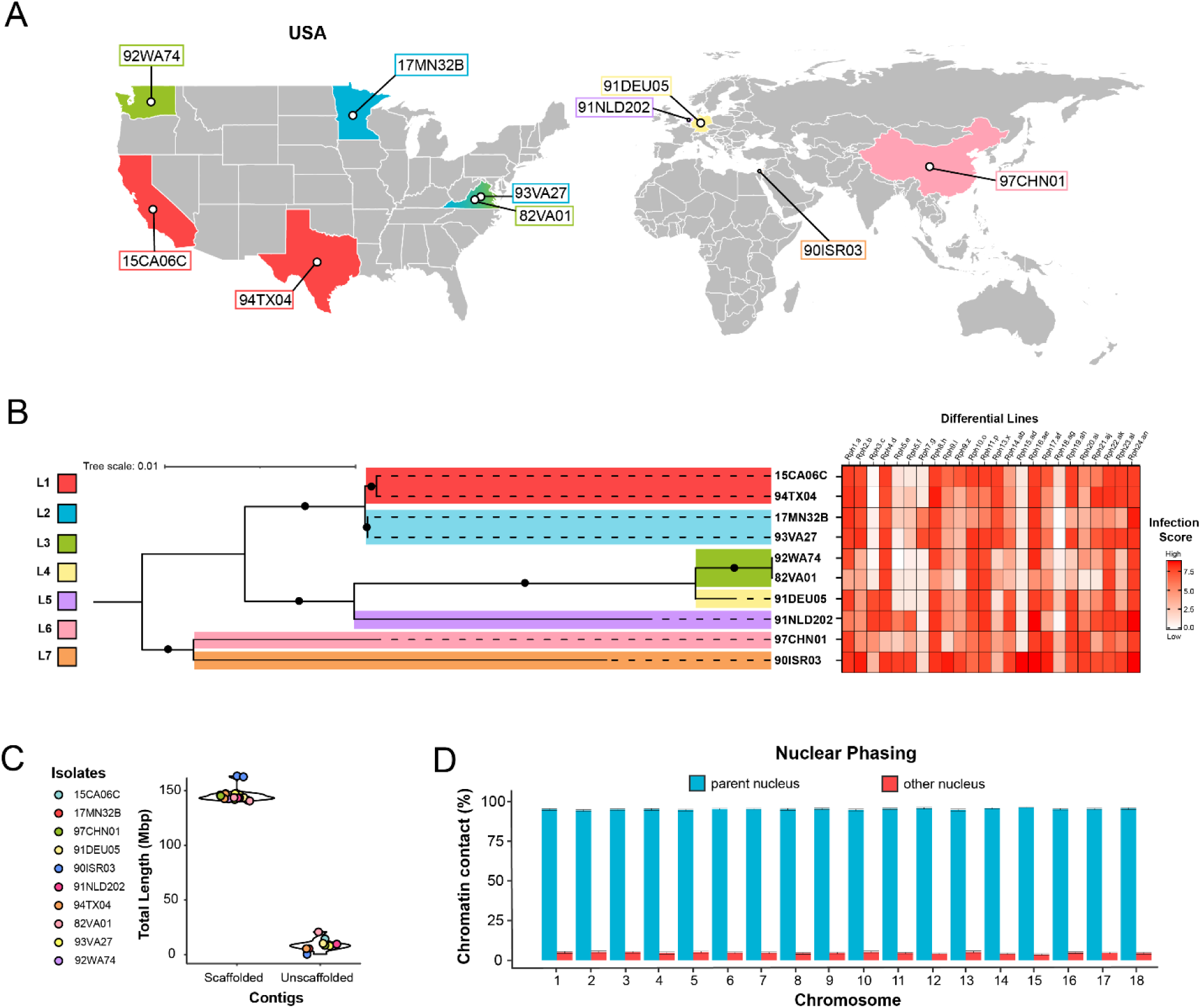
A) Geographic origin of isolates (US state or country) are marked and color-coded by lineage as in panel 1A. B) Maximum likelihood tree showing the phylogenetic relationship among the ten *P. hordei* diploid genomes. All genomes were aligned to the diploid 17MN32B genome. Branches supported by ≥80% of 500 bootstraps are represented by a black circle in the middle. The seven identified clonal groups (lineages) are color-coded. Infection scores (from 0-9; where 0=a completely incompatible reaction with no symptoms and 9=complete compatibility with extensive sporulation) on differential lines containing the leaf rust resistance genes *Rph1-24*. C) Violin plot of total scaffolded contig lengths (bp) and total unscaffolded contig lengths (bp) in haploid (haplotype-phased) *P. hordei* assemblies. D) The mean (±SD) proportion of *trans* Hi-C contacts (%) of each chromosome to both the parent nucleus and the other nucleus in the dikaryon across all nuclear haplotypes.

### Genome sequencing and assembly

High molecular weight DNA was extracted from ∼450 mg purified urediniospores for long read PacBio HiFi sequencing as described by Li et al. (2019). Samples were sequenced at 30-50X depth at the University of Minnesota Genomics Center (UMGC, Oakdale, MN) in pooled libraries of 5 samples over two SMRT cells on the Sequel II platform. For Hi-C sequencing, 300 mg fungal spores of each isolate were incubated in 1% formaldehyde for 20 min at room temperature with intermittent vortexing to crosslink DNA, followed by the addition of glycine (to 125mM final concentration) and an additional incubation of 15 min. Spores were washed twice with distilled water and collected via filtration using cell filters. Then, the spores were ground to a fine powder in liquid nitrogen and sent to Phase Genomics (Seattle, WA, USA) on dry ice for preparation of sequencing libraries. The Phase Genomics Proximo Fungal kit version 4.5 was used for preparation of an 150bp insert paired-end library. Libraries were sequenced on the Illumina NovaSeq X Plus platform by Azenta Life Sciences (South Plainfield, NJ, USA).

GenomeScope 2.0 (Ranallo-Benavidez et al., 2020) was used to generate a histogram of *k*-mers from PacBio HiFi reads and infer ploidy (signs of contamination), level of heterozygosity, and genome size. Hifiasm v0.19.9-r616 (Cheng et al., 2021) was used in Hi-C integration mode to assemble initial haplotype-phased genomes using the corresponding PacBio HiFi and Hi-C sequencing reads. Assembly quality was assessed using BUSCO v3 (Simão et al., 2015) (-l basidiomycota_odb9 -sp coprinus) and GenomeScope v2.0 (Ranallo-Benavidez et al., 2020). Contig coverage was evaluated by mapping PacBio HiFi reads to the hifiasm assembly (minimap2 v2.26-r1175 (H. Li, 2018)) and using the custom script Coverage.R (https://github.com/JanaSperschneider/GenomeAssemblyTools/CollapsedGenomicRegions/ Coverage.R). Low coverage contigs were removed after determining thresholds from whole-genome coverage distribution plots. Contaminant and mitochondrial contigs were removed after BLASTn searches of the NCBI nucleotide (-e-value 1e-5 -perc_identity 75) and mitochondrial databases (-perc_identity 90.0). Phasing was assessed by running NuclearPhaser (https://github.com/JanaSperschneider/NuclearPhaser,(Duan et al., 2022)) with MAPQ=30 and necessary phase switches >400kb were performed. Scaffolds were generated with SALSA (Ghurye et al., 2017) and a synteny-guided *de novo* assembly of isolate 17MN32B was performed by aligning scaffolds to the individual haplotypes of *P. graminis* f. sp. *tritici* Pgt21-0 (hapA, hapB) using D-GENIES (Cabanettes & Klopp, 2018). A custom scaffolding script was used to build full chromosomes and assess the presence of telomeres (https://github.com/JanaSperschneider/FindTelomeres). Manual curation of chromosomes was performed by building chromatin contact matrices with HiC-Pro v3.1.0 (Servant et al., 2015) and assessing chromatin contact plots using hicexplorer (Wolff et al., 2020). The chromosomes for the remaining *P. hordei* assemblies were scaffolded using a combination of scaffolds generated from SALSA (Ghurye et al., 2019) scaffolder and also by aligning to the phased haplotypes of isolate 17MN32B using D-GENIES and inspecting the order of contigs in the plot. Chromatin contact plots were used to curate the final chromosomes as described above. The quality of the final chromosome-level haplotype-phased assemblies were assessed again using BUSCO v3 (Simão et al., 2015) (-l basidiomycota_odb9 -sp coprinus). HiCanu (Nurk et al., 2020) assemblies were also performed using the PacBio HiFi data.

### Nanopore sequencing

Oxford Nanopore Technologies (ONT) sequencing was performed for isolates 15CA06C, 92WA74 and 90ISR03 using HMW DNA extracted using the method described above. Sequencing libraries were prepared using the short read eliminator kit (Cat No: EXP-SFE001) and ligation sequencing kit V14 (Cat No: SQK-LSK114). One to two libraries per isolate were loaded onto a flow cell (Cat No: FLO-MN114) for sequencing on the MinION Mk1B (Cat No: MIN-101B) sequencer (Oxford Nanopore Technologies, Oxford, UK). The quality of reads was assessed using FastQC (Andrews, 2010). Dorado (https://github.com/nanoporetech/dorado, Oxford Nanopore Technologies) was used to basecall pod5 reads. Fastq reads were aligned to the diploid genome assembly using minimap2 (Li, 2018) for inspection.

### RNA sequencing

RNA sequencing was performed for germinated urediniospores of isolate 82VA01 and also barley leaves infected with the same isolate, in triplicate. Fresh 82VA01 spores were initially dusted over germination solution (Barnes & Szabo, 2007). After 10h, the mat of germinated urediniospores was harvested using a spatula. The susceptible barley cultivar Robust (PI 476976) was inoculated with urediniospores of 82VA01 according to the methods described above, and then infected leaf tissue samples were harvested at 5 and 7 days post-infection (DPI). All samples were flash-frozen upon harvest and ground in liquid nitrogen. RNA was extracted using the Qiagen RNeasy Plant Mini Kit (Cat No: 74904, QIAGEN, Aarhus, Denmark). All samples were sequenced at UMGC on the Illumina NextSeq P3 platform after preparation of a 150bp paired-end TruSeq stranded mRNA library. The quality of RNAseq data was assessed through STAR aligner read mappings to the phased 82VA01 reference (Dobin et al., 2013).

### Genome annotation

For gene prediction, the haplotype-phased *P. hordei* genomes were soft-masked with RepeatMasker v4.0.5 (with options -s -nolow -lib) (Tarailo-Graovac & Chen, 2009) using the RepeatModeler v2.0.1 libraries with unknown repeats removed (Flynn et al., 2020). Trimmomatic (v0.39) (Bolger et al., 2014) was used to trim *P. hordei* isolate 82VA01 paired-end Illumina RNAseq reads (with options ILLUMINACLIP:TruSeq3-PE.fa:2:30:10:2:True LEADING:3 TRAILING:3 MINLEN:36) and unpaired reads were removed. Trimmed RNAseq reads of isolate 82VA01 were mapped to the diploid assemblies (including individual haplotypes and unplaced contigs) using HiSat2 v.2.1.0 (Kim et al., 2019) (--max-intronlen 3000 --dta --no-unal --rna-strandedness RF). Read mappings were sorted, indexed and merged (samtools v1.16.1, (Danecek et al., 2021)) and Trinity v2.15.0 (Bankar et al., 2015) was used in genome-guided assembly mode with options (--genome_guided_bam --genome_guided_max_intron 3000 – jaccard_clip) to generate transcripts (fasta). StringTie v1.3.4 (-s1 -m50 –rf) (Pertea et al., 2015) was used to assemble transcripts for each sample from the HiSat2 alignments and merge them into separate pools of transcripts from germinated urediniospores and barley leaves infected with *P. hordei*. CodingQuarry (Testa et al., 2015) was used to predict genes in the diploid assemblies and was run on the infected barley leaf transcripts and germinated urediniospore transcripts separately (filtering performed as described in Sperschneider *et al*. 2022).

Funannotate v. 1.7.4 (Palmer & Stajich, 2020) training was run on preassembled Trinity transcripts (with options –stranded RF –no-trimmomatic –jaccard_clip). Funannotate predict was run on the soft-masked genomes using CodingQuarry (-other_gff with weight of CodingQuarry infection gene predictions set to 20 and weight of spore gene predictions set to 2), and pucciniomycotina EST evidence downloaded from JGI MycoCosm (--ploidy 2 –optimize_augustus –busco_seed_species ustilago –weights pasa:10 codingquarry:0). Funannotate update was then run to add UTRs and refine predictions. To manually capture additional small secreted proteins (putative effectors), open reading frame (ORF) prediction was performed using TransDecoder v5.5.0 (Haas & Papanicolaou, 2017) on the StringTie infection transcripts (TransDecoder.LongOrfs -m50 and TransDecoder.Predict –single_best_only). We filtered for complete ORFs (with ‘start’ and ‘stop’ codons) with a signal peptide (SignalP 4.1 -u 0.34 -U 0.34) and no transmembrane domains outside the N-terminal signal peptide region (TMHMM 2.0) (Möller et al., 2001; Nielsen, 2017). Genes encoding secreted proteins were added to the annotation file using agat_sp_fix_overlaping_genes.pl (https://github.com/NBISweden/AGAT), which creates isoforms for genes with overlapping coding sequence. Genes that were 50 bp or longer in length were retained (agat_sp_filter_gene_by_length.pl with options –size 50 –test “>”). Only the longest isoform was retained for each gene (agat_sp_keep_longest_isoform.pl). The completeness of annotation was assessed using BUSCO v3 (-m prot -l basidiomycota_odb9 -sp coprinus) (Simão et al., 2015). InterProScan v5.23 (Jones et al., 2014) was run to predict the function of annotated proteins. Functional annotation of gene models was performed using funannotate annotate.

### Pan-genome saturation analysis

To assess pan-genome completeness, orthogroup clustering results from OrthoFinder v2.5.4 (Emms & Kelly, 2019) were used to construct a binary presence/absence matrix across 13 unique *P. hordei* haplotypes. This was used as the input for R package micropan v2.1 (Snipen & Liland, 2015). Rarefaction was performed with 1,000 random permutations of haplotype order to estimate the accumulation of orthogroups with increasing haplotype sampling. Mean values across permutations were plotted for each haplotype value (1-13) for the pan-genome (cumulative number of orthogroups observed), core genome (present in all haplotypes) and unique genome (present in only one haplotype). micropan function heaps was used to estimate the Heaps’ Law decay parameter α. The Chao lower bound (Chao, 1987) was estimated using micropan function chao.

### Comparative genomics

Funannotate compare was run on the annotated gene models. OrthoFinder v2.5.4 (Emms & Kelly, 2019) was used to identify gene orthologs and orthogroups, together with their phylogenetic relationships and gene duplication events. EffectorP v3.0 (Sperschneider & Dodds, 2022) was run on proteins with identified secretion signals (SignalP v4.1 -u 0.34 -U 0.34) to predict secreted cytoplasmic and apoplastic effector proteins from each isolate. AntiSMASH v7.0 fungal version (Blin et al., 2023) was run to predict secondary metabolite biosynthesis gene clusters in each phased genome.

### Mitochondrial genome assembly

The MitoHiFi pipeline (https://github.com/marcelauliano/MitoHiFi.git (Uliano-Silva et al., 2023)) was used for mitochondrial genome assembly with the unfiltered hifiasm contigs for isolate 15CA06C using the wheat leaf rust pathogen *Puccinia triticina* isolate HnZU18-3 (MN004749.1) mitogenome contigs and GenBank entry as references. The pipeline filters input contigs that align to the reference sequence to select the longest contig containing orthologous mitochondrial genes, which is then circularized. OGDRAW (Greiner et al., 2019) was used to draw a mitochondrial genome map.

### Repeat annotation

Haplotype-phased chromosome-level genome assemblies for other dikaryotic rust species were downloaded from NCBI or elsewhere to compare repeat content (**Table S22**). De novo repeats were predicted with RepeatModeler version 2.0.1 (Flynn et al., 2020) utilizing the option - LTRStruct. RepeatMasker (-s -engine ncbi) (Tarailo-Graovac & Chen, 2009) was run with the RepeatModeler library to obtain repetitive element statistics.

### Mating type allele analyses

Protein sequences for STE3.2-2, STE3.2-3, mfa2, mfa3, bW-HD and bE-HD from *Puccinia graminis* f. sp. *tritici* CRL 75-36-700-3 (Cuomo et al., 2017) were used for tblastn searches (NCBI toolkit v. 25.2.0) to locate the corresponding gene homologs in *P. hordei*. Coding sequences were extracted in each isolate. The NGPhylogeny.fr pipeline (Lemoine et al., 2019) was used to integrate multiple tools for alignment of the *STE3.2-2*, *STE3.2-3*, *bW-HD1* and *bW-HD2* alleles to their respective homologous sequences (MAFFT, (Katoh & Standley, 2013)) followed by alignment curation (BMGE (Criscuolo & Gribaldo, 2010)) and generation of a maximum likelihood tree (PhyML, (Guindon et al., 2010)) with 1000 bootstraps. *K*-mer containment of *bW-HD1* and *bE-HD2* alleles within available Illumina reads was performed using mash (Ondov et al., 2016).

### Phylogenetic analysis

Minimap2 v2.26-r1175 (Li, 2018) was used with option -ax asm5 to align diploid assemblies to the phased 17MN32B assembly, and all haploid assemblies (haplotypes) to the 15CA06C hapA assembly. Samtools v1.21 (Danecek et al., 2021) was used to sort and index the resulting bam files. Bcftools v1.2 (Danecek et al., 2021) mpileup and call were used to call SNPs and indels (-50bp) in the alignments against the indexed reference sequence. Heterozygote calls between the haploid genomes were converted to missing. For both diploid and haploid vcfs, vcftools (Danecek et al., 2011) was used to keep genotyping sites with ≤ 10% missing data. Vcf files were converted to phylip files using the script: https://github.com/edgardomortiz/vcf2phylip. RAxML v8.2.11 (Stamatakis, 2014) was used to generate a single phylogenetic tree (-f b -z -t -m GTRCAT) comprised of 500 bootstrap trees (-f a -# 500 -m GTRCAT) and a single maximum-likelihood tree (-D).

### Pairwise sequence comparisons

D-GENIES (Cabanettes & Klopp, 2018) pairwise alignments of haplotype-phased assemblies was performed to aid scaffolding and visualize translocations. Haplotypes were aligned with nucmer from the MUMmer package (Marçais et al., 2018) and divergence was calculated using dnadiff. Mash v2.3 (Ondov et al., 2016) was used to measure containment of diploid assemblies, haplotype-phased assemblies, and mating type alleles in raw genomic sequencing reads (using -s 10,000 for assemblies and -s 1000 for alleles). For visualization of syntenic blocks, minimap2 (H. Li, 2018) with option -ax asm5 was used to perform whole-genome alignment between haplotype-phased genome assemblies. Syri (Goel et al., 2019) was performed on each alignment file to annotate syntenic blocks and structural rearrangements. The resulting annotations were visualized using plotsr (Goel & Schneeberger, 2022).

## Results

### Sequenced *Puccinia hordei* isolates exhibited diverse virulence profiles

To capture maximal genetic diversity, *P. hordei* isolates were selected for sequencing based on 1) geographic origin, 2) avirulence phenotypes on barley leaf rust differential lines (Martin et al., 2020), 3) year of sampling and 4) previous genotyping with AFLP markers. AFLP genotyping of a diverse collection of 45 *Ph* isolates identified six molecular groups (MGs) (Sun et al., 2007). Ten *Ph* isolates spanning sampling years 1982-2017 were selected for PacBio HiFi and Hi-C sequencing: 15CA06C (California, USA, 2015), 17MN32B (Minnesota, USA, 2017), 97CHN01 (MGIII, China, 1997), 91DEU05 (MGV-1, Germany, 1991), 90ISR03 (MGII, Israel, 1990), 91NLD202 (MGIV-4, Netherlands, 1991), 94TX04 (MGV-2, Texas, USA, 1994), 82VA01 (Virginia, USA, 1982), 93VA27 (MGIV, Virginia, USA, 1993) and 92WA74 (MGIV-2, Washington, USA, 1992) (**Figure 1A, Table S1**). Isolates exhibited a broad range of phenotypes on the barley leaf rust differential lines containing the *R* genes *Rph1*-*Rph24* (**Figures 1B, S1, Table S2**). Notably, 90ISR03 from Israel was the most broadly virulent isolate and is the only reported isolate with virulence on *Rph15* to-date (Sun et al., 2007) (**Figures 1B, S1, Tables S2, S3**). The isolate with the narrowest virulence spectrum was from the US (82VA01). All isolates were virulent on *Rph8*, *Rph11*, *Rph16*, *Rph22* and *Rph24*, whilst *Rph18* provided resistance against all isolates.

### Establishment of a *Puccinia hordei* nuclear haplotype collection

Nuclear haplotype-phased reference genomes were assembled using hifiasm (Cheng et al., 2021) in Hi-C integration mode (**Table S5**). For isolate 17MN32B, contigs were initially scaffolded using a combination of SALSA and synteny with *P. graminis* f. sp. *tritici* isolate *Pgt*21-0 chromosomes, with Hi-C contact maps used to confirm correct chromosome construction. Chromosomes were numbered based on synteny with *P. graminis* f. sp. *tritici*, consistent with practice for other *Puccinia* species (Henningsen et al., 2024; Liang et al., 2023; Sperschneider et al., 2023; Yu et al., 2025). The remaining *P. hordei* isolates were scaffolded using a combination of SALSA and synteny with the 17MN32B chromosomes, using chromatin contact maps to curate the final chromosomes. For each chromosome, an average of 93.5% *trans* Hi-C contacts and 97.2% *cis* and *trans* contacts occurred within its designated nucleus (**Figure 1D, Table S4**). This indicated accurate phasing and establishment of nuclear haplotypes and was supported by the Hi-C chromatin contact maps. Phase switches of >400kb were only detected in the assemblies from isolates 90ISR03 (x1), 82VA01 (x1) and 93VA27 (x2) and all were corrected prior to scaffolding. Nine of the ten isolates were scaffolded into 18 chromosomes with 10-56 gaps per haplotype. 90ISR03 had an extra chromosome-like scaffold of ∼6.2Mbp in hap6 with a centromere-like region and two telomeres that could not be scaffolded on core chromosomes (designated chromosome 19). Unscaffolded contigs ranged from 300kbp-20.66Mbp (**Figure 1C**, **Table S5**).

The genetic relationship of *P. hordei* isolates was determined by aligning the diploid assemblies to the haplotype-phased 17MN32B assembly as a reference (**Figure 1B**). Seven distinct lineages were found based on phylogenetic analysis, with the six US isolates falling into three pairs representing clonal lineages. These lineages were reflected in the phenotyping data (**Figure 1B**). A circularized mitochondrial genome (39 genes) was also assembled for isolate 15CA06C using the MitoHiFi pipeline (**Figure S2**). Alignment of PacBio HiFi reads from the nine remaining isolates to this assembly revealed that they shared identical mitochondrial genomes.

Genome completeness and integrity were assessed through multiple approaches. The BUSCO v3 completeness score was consistent across haplotypes with an average of 94.8% and 1.7% duplication, which was comparable to previous phased assemblies of other *Puccinia* species (Henningsen et al., 2024; Li et al., 2019; Sperschneider et al., 2023; Wang et al., 2024). Four of ten diploid and ten of 20 haploid assemblies were complete telomere-to-telomere with 17 of 20 haploid assemblies harboring 35+ telomeres (**Table S6**). This represents the most structurally complete set of phased *P. hordei* genomes to date. Pairwise nucleotide alignments of 17MN32B haplotypes to haplotype-phased assemblies from other *Puccinia* species revealed that *P. hordei* had closest sequence identity to *P. coronata* f. sp. *avenae* (oat crown rust pathogen, isolate Pca203) (86.59-86.88%), followed by *P. triticina* (wheat leaf rust pathogen, isolate 20QLD87) (85.22-85.25%), *P. graminis* f. sp. *tritici* (wheat stem rust pathogen, isolate ETH2013-1) (85.14-85.32%) and *P. striiformis* f. sp. *tritici* (wheat stripe rust pathogen, isolate AZ2) (84.93-85.36%) (**Table S7**). The average haplotype size of *P. hordei* was 146.13 Mb, which was consistent with the phased haplotype sizes obtained by Yu et al. (2025) (**Figure 1C**, **Table S6**). Nine isolates had haplotype sizes ranging from 140.69-147.41Mbp, with isolate 90ISR03 harboring larger haplotypes of 164.8 and 161.4Mbp (**Table S6**).

### Somatic hybridization in *Puccinia hordei* was evidenced by shared nuclear haplotypes

Since nuclear exchange has been demonstrated in multiple *Puccinia* species (Henningsen et al., 2024; Holden et al., 2025; Li et al., 2019; Sperschneider et al., 2023), leading to distinct lineages sharing nuclei, we investigated the containment of nuclear haplotypes in this *P. hordei* haplotype collection and available Illumina data for other isolates of the pathogen. Pairwise nucleotide identity of *P. hordei* nuclear haplotypes mostly ranged from 98.77-99.98%, with the haplotypes from 90ISR03 being the most diverged, with only 98.48 to 98.53% identity to haplotypes from other isolates (**Table S8**). Both pairwise alignment and containment of haplotypes in the PacBio HiFi reads (and within haplotypes) revealed the presence of 13 distinct haplotypes from 20 nuclear assemblies (**Figure 2A**, **Tables S8, S9**). Due to these similarities, we named the 13 distinct nuclear haplotypes hap1-hap13 (**Table S10**). Three clonal lineages were present within the US isolates, each represented by two isolates, as reflected in phylogenetic analyses of diploid (**Figure 1B**) and haploid assemblies (**Figures 2A, 2B, S3**). Two of the clonal US lineages (L1, L2) shared hap2, indicating at least one somatic hybridization event in the emergence of these lineages (**Figures 2A, B**). Within the three clonal lineages, shared nuclear haplotypes had 4-8,000 SNPs between them, whereas divergent haplotypes had 600-800,000 SNPs (**Figure 2C**). Hap2 was also contained in the Illumina reads of Australian isolate Ph560 (Chen et al., 2019) and its mutational derivatives (**Figure 2B**, **Table S11)**.

**Figure 2.**
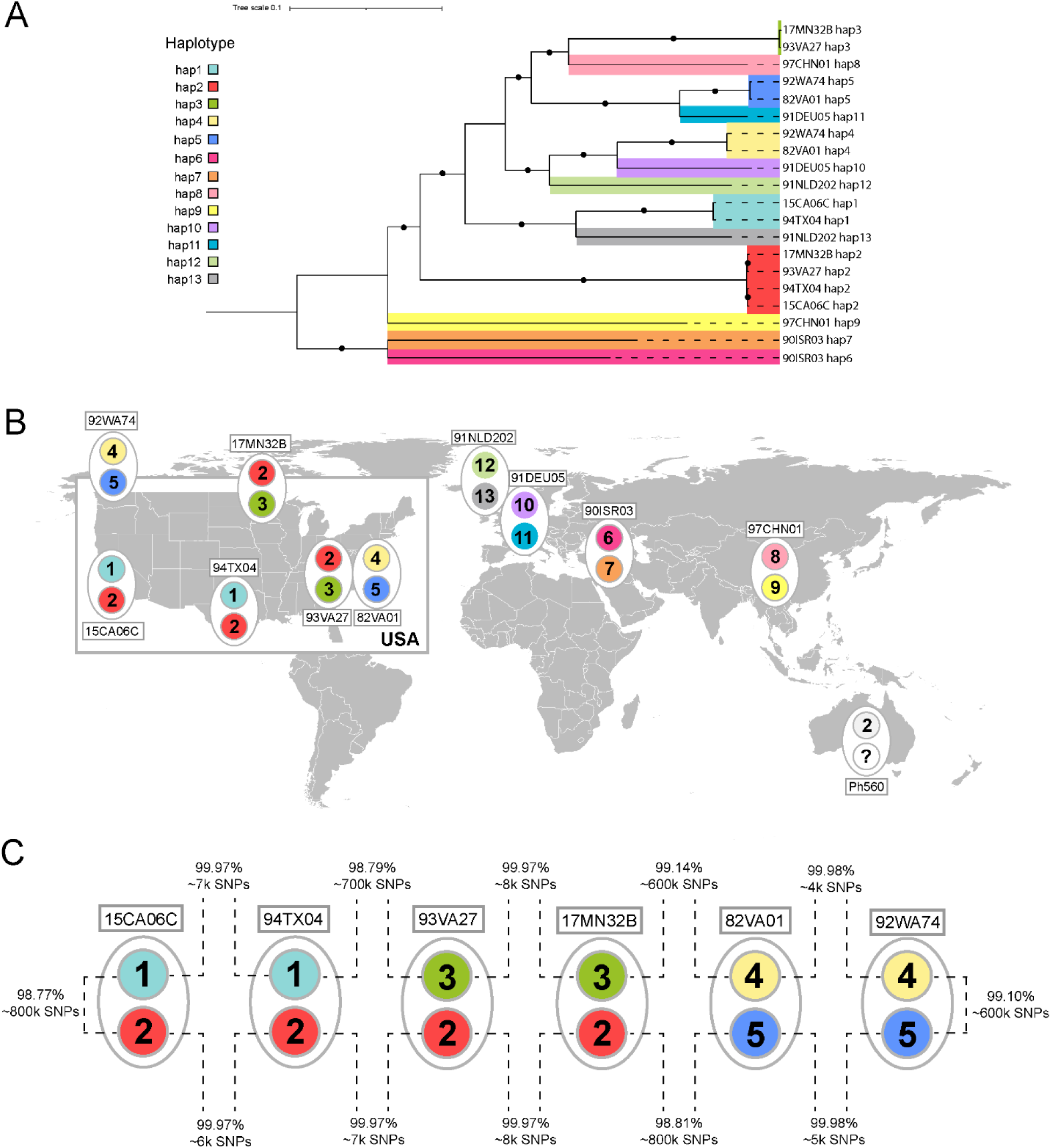
A) Maximum likelihood tree showing the phylogenetic relationship between the 20 *P. hordei* haploid genomes, color-coded by nuclear haplotype identity (hap1-hap13), after alignment to 15CA06C hap1. Branches supported by ≥80% of 500 bootstraps are represented by a black circle in the middle. B) Geographic origin and haplotype identity of *P. hordei* isolates, inferred by whole-genome pairwise alignment (bold colors) and postulated via *k*-mer containment of phased nuclear haplotypes in Illumina reads (light grey). C) Nucleotide identity (%) and number of SNPs between nuclear haplotypes in the three clonal US lineages, derived from mummer (nucmer) pairwise alignments of haploid assemblies.

### High genome plasticity was observed as extensive structural rearrangements

Compared to other *Puccinia* species (Henningsen et al., 2024; Li et al., 2019; Sperschneider et al., 2023; Wang et al., 2024), *P. hordei* exhibited a much larger range of chromosome sizes (**Figure 3B, Table S12**). The most variable chromosome lengths were chr1 (6.19-12.88Mb), chr2 (9.79-14.17Mb), chr8 (4.93-9.06Mb) and chr13 (7.09-11.65Mb). The extremes in chromosome sizes could largely be explained by reciprocal translocations in some isolates and/or haplotypes including: between chr3 and chr8 in hap1 of 15CA06C; chr1 and chr13 in hap2 of 17MN32B; and chr1 and chr8 in hap4 of 92WA74 and hap4 of 82VA01 (**Figures 3B, 3C, S4**). Although the chr1-chr8 translocation was conserved between the shared hap4 in clonal isolates 92WA74 and 82VA01, the chr3-chr8 translocation in 15CA06C hap1 was not present in hap1 of the clonal isolate 94TX04. Likewise, the chr1-chr13 translocation in 17MN32B hap2 was not present in hap2 of the clonal isolate 93VA27. To differentiate between real translocations and artefacts of mis-assembly due to repetitive regions, we accumulated several independent sources of evidence to support the translocations. First, we inspected Hi-C chromatin contact maps and found continuous contact across these chromosomes (**Figure S5**). Then we generated independent *de novo* assemblies for these isolates using the HiCanu assembler and found that HiCanu contigs also supported the putative breakpoints (**Figure S5**). Lastly, PacBio HiFi reads were mapped to the phased assemblies and reads were identified that spanned the chromosome breakpoints (**Figure S6**). As further support, we obtained Oxford Nanopore long reads for isolates 15CA06C and 92WA74. We mapped these reads to the putative translocated chromosomes and found continuous coverage across the full length of the chromosomes (**Figure S7, Table S13**). A number of additional structural variants were identified including inversions and duplications (**Figure 3A**).

**Figure 3.**
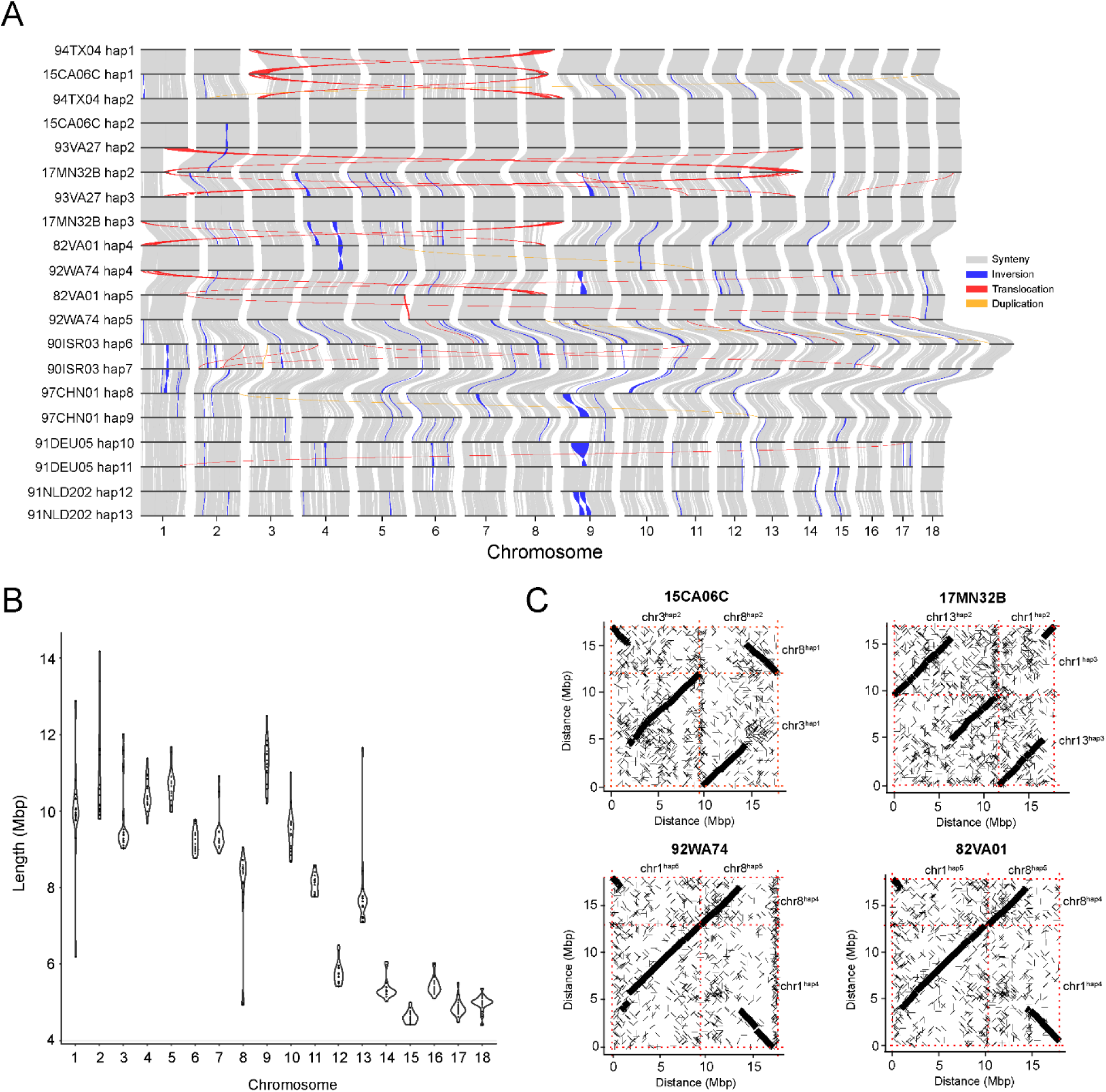
A) Synteny among chromosomes 1-18 in the twenty nuclear-phased haplotypes of ten *Puccinia hordei* isolates. B) Chromosome lengths (Mbp) in chromosomes 1-18 of the 20 nuclear-phased haplotypes of *P. hordei*. C) Dotplots showing pairwise alignments (D-Genies) between the structurally-rearranged chromosomes of 15CA06C hap1, 17MN32B hap2, 92WA74 hap4 and 82VA01 hap4 and the homologous chromosomes in the other nuclear haplotype.

### Identification of a supernumerary chromosome in one haplotype of 90ISR03

The isolate 90ISR03 had consistently larger chromosomes than the other isolates (ranking either first or second largest) (**Figure 3B, Table S12**). 90ISR03 also had an extra chromosome-like scaffold of 6.2Mbp in hap6 which we provisionally designated as chr19 (**Table S5, Figure S8**). The supernumerary scaffold had both forward and reverse telomeres, and a clear centromeric-like region (alpha-satellite repeats) when inspecting HiC chromatin contact maps (**Figure S9**). Chromatin contact data suggested that the scaffold was phased correctly, was only present in hap6, and that its two constituent contigs (∼4.8Mbp and ∼1.4Mbp) could not be scaffolded elsewhere (**Table S14**, **Figure S10**). The first ∼1.4Mbp of chr19 sequence is also present but inverted (reverse complemented) in the final 1.4Mbp (4.8-6.2Mbp). These terminal sequences have 99.99% sequence identity with only eight SNPs detected in pairwise alignment. Inspection of the PacBio HiFi read mappings demonstrated coverage across the entire 6.2Mbp chromosome, including over the boundaries of the duplicated regions at ∼1.4Mbp and ∼4.8Mbp, the latter of which is also the boundary between the two contigs in the scaffold (**Figure S11**). As further support for the additional scaffold, we generated Oxford Nanopore long reads for this isolate and found continuous coverage of reads mapped across the scaffold (**Table S13, Figure S12**). PacBio HiFi or Nanopore reads mapping to the ends of the scaffold did not map elsewhere. A HiCanu assembly of 90ISR03 contained three contigs of ∼3.74Mbp, ∼1.05Mbp and ∼50kbp that aligned to chr19, giving further support for its existence. The 3.74Mbp contig aligned to the first ∼3.74Mbp of chr19 with 99.99% nucleotide identity and also aligned from ∼4.8Mbp to the end, suggesting that the duplicated terminal region had collapsed in this assembly (**Figures S13, S14**). The other smaller HiCanu contigs of 1.05Mbp and 50kbp mapped to regions in the middle of chr19 with 99.99% nucleotide identity (3.78-4.83Mbp, and 3.73-3.78Mbp, respectively) (**Figures S13, S14**).

The chr19 scaffold was not contained in other *P. hordei* pan-genome isolates as assessed through pairwise alignment and *k*-mer containment (**Table S15**). BLASTn alignment of the chr19 sequence revealed closest matches with *P. hordei* chromosomal sequence. Significant hits were found to all chromosomes with the longest significant matching region being ∼13,000 bp on chr5. The PacBio HiFi read coverage for chr19 was comparable to core chromosomes (**Table S16**). A total of 522 genes were annotated on chr19. None of these were BUSCO (n=1335) genes. Gene density for chr19 was one gene per 11,918 bp which was lower than the core chromosome average of one gene per 8,807 bp (range of one per 7,520-9,792 bp) (**Figure S8**). The repeat coverage of chr19 was 85.83% which was higher than chr9^hap6^ (81.89%) and chr9^hap7^ (81.89%) which are known to have large repetitive regions encoding ribosomal RNA genes (higher repeat content than other core chromosomes) (Yu et al., 2025). A total of 25 predicted secreted proteins were annotated on chr19 and three of these were predicted effectors (EffectorP). No secondary metabolite clusters were predicted on chr19.

### Characterization of mating type loci in the *P. hordei* haplotype collection

Similar to other *Puccinia* species (Henningsen et al., 2024; Li et al., 2019; Sperschneider et al., 2023), *P. hordei* harbors a bi-allelic mating type locus *a* on chromosome 9 comprised of pheromone receptor *STE3.2-2* or *STE3.2-3* and a linked mating factor *a* pheromone precursor (*mfa2* or *mfa3*, respectively) (**Figure 4A**). This locus was universally heterozygous between the *P. hordei* dikaryotic nuclei and no genic variation was found among isolates. The mating type *a* locus is embedded in a large repeat-rich region on chromosome 9 (in a homologous region in *P. coronata* f. sp. *avenae* (Henningsen et al., 2024), *P. graminis* f. sp. *tritici* (Li et al., 2019; Spanner et al., 2025) and *P. triticina* (Sperschneider et al., 2023), **Figure 4A**). The two distinct alleles are associated with repeat regions of different sizes, as reflected by the total length of chromosome 9 (**Figure 4C, D**). The *STE3.2-2* allele was in head-to-head orientation with *mfa2* with a gap of ∼900bp between start codons, suggesting a shared promoter region (**Figure 4A**). The *STE3.2-3* allele was upstream of *mfa3* in tail-to-tail orientation with a variable intergenic region between isolates (ranging from 88 to -234kb, with an average gap of 163 kb). The *b* locus was composed of two tightly-linked homeodomain transcription factors (*bW*-*HD1* and *bE-HD2*) on chromosome 4 (**Figure 4B**). A total of 10 distinct alleles were found for the two *HD* genes (*bWbE* alleles) (**Tables S17, S18**). Shared nuclei harbored identical *bWbE* alleles. The *bW1* and *bE1* alleles were shared by four distinct haplotypes (hap1, hap4, hap10 and hap12). The *bW2* and *bE2* alleles found in hap2-containing isolates were contained in the reads of Australian isolate Ph560 and the clonal isolates Ph584, Ph608, Ph612, Ph626, as expected (**Table S19**).

**Figure 4.**
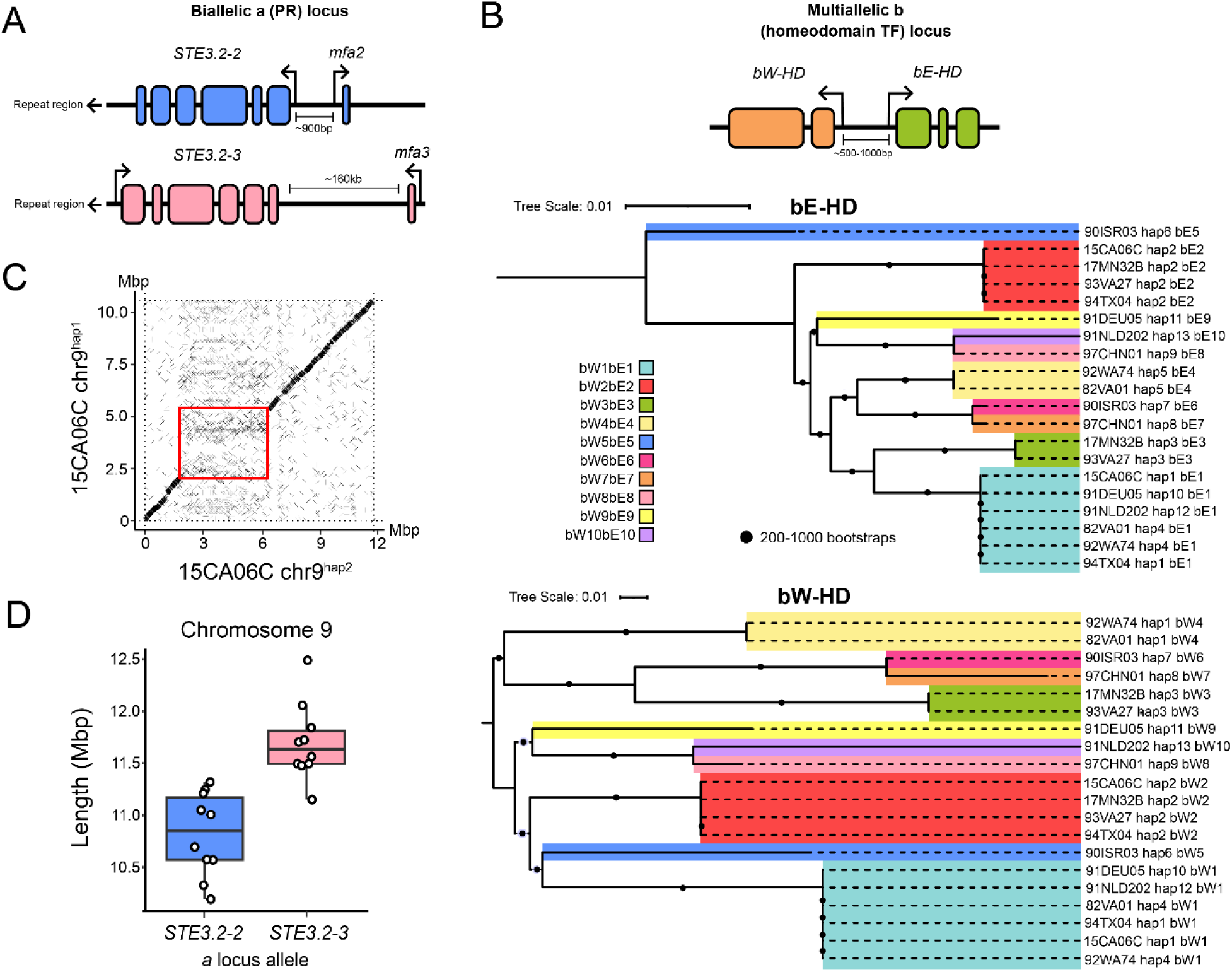
A) The biallelic mating type *a* locus with the genomic context of pheromone receptor genes *STE3.2-2* and *STE3.2-3* and linked pheromone precursor genes *mfa2* and *mfa3* depicted. B) The top panel depicts the multiallelic *b* mating type locus with the genomic context of homeodomain transcription factor genes *bW-HD1* and *bE-HD2*. A maximum likelihood tree with 1000 bootstraps demonstrating the phylogenetic relationships between the 10 identified alleles amongst the 20 *P. hordei* haplotypes for the *bWbE* locus. Branches supported by ≥80% of bootstraps are represented by a black circle in the middle. C) Pairwise alignment (D-Genies) between chromosome 9^hap1^ and chromosome 9^hap2^ of isolate 15CA06C, demonstrating the location of the large repeat block on both chromosomes and positions of *STE3.2-2*/*STE3.2-3* mating type genes. D) Length of chromosome 9 with either the *STE3.2-2* or *STE3.2-3 a* locus associated with repeat regions of distinct sizes.

### Repeat content of *Puccinia hordei* genomes

The repeat content of *P. hordei* was 69.6-70.8% among nine isolates, with a higher repeat content of ∼72% for isolate 90ISR03 (**Tables S6, S20**). The total length of repetitive sequence for *P. hordei* 17MN32B hap2 was ∼104Mbp, which was higher than for *P. triticina* 19NSW04 hapB (∼71Mbp), *P. coronata* f. sp. *avenae* 203 hapA (∼54Mbp), *P. graminis* f. sp. *tritici* ETH2013-1 hap4 (∼36Mbp), *P. striiformis* f. sp. *tritici* AZ2 hap1 (∼28Mbp) and *P. silphii* (∼12Mbp). Since the non-repetitive part of the genome is consistent in size between these species (except microcyclic *P. silphii*), we can attribute the larger size of the *P. hordei* genome (∼145 Mbp), and indeed the size variation between the other *Puccinia* species, predominantly to repeat element expansion (**Figure 5A, Table S22**). *P. hordei* had notably longer total sequences of long terminal repeat (LTR) retroelements Gypsy/DIRS1/Ty1/Copia, DNA transposons, non-LTR retroelements and rolling circle transposons when compared to representative chromosome-level haplotype-phased genomes from other *Puccinia* species (**Figure 5B**), suggesting that expansion has occurred in all major classes of repeat elements. When looking at intraspecific variation in repeat content, isolate 90ISR03 had a larger genome than the other *Ph* isolates in the haplotype collection; this was partially due to harboring an extra chromosome in one haplotype but is also attributed to a higher repeat content when assessing the core chromosomes 1-18, as presented in **Figure 5C** (**Tables S20**, **S21**). Notably, there was more sequence assigned to Ty1/Copia/Gypsy/DIRS1 LTR elements and Tc1-IS630-Pogo/Tourist/Harbinger DNA transposons in 90ISR03 compared to the other isolates (**Figure 5C**).

**Figure 5.**
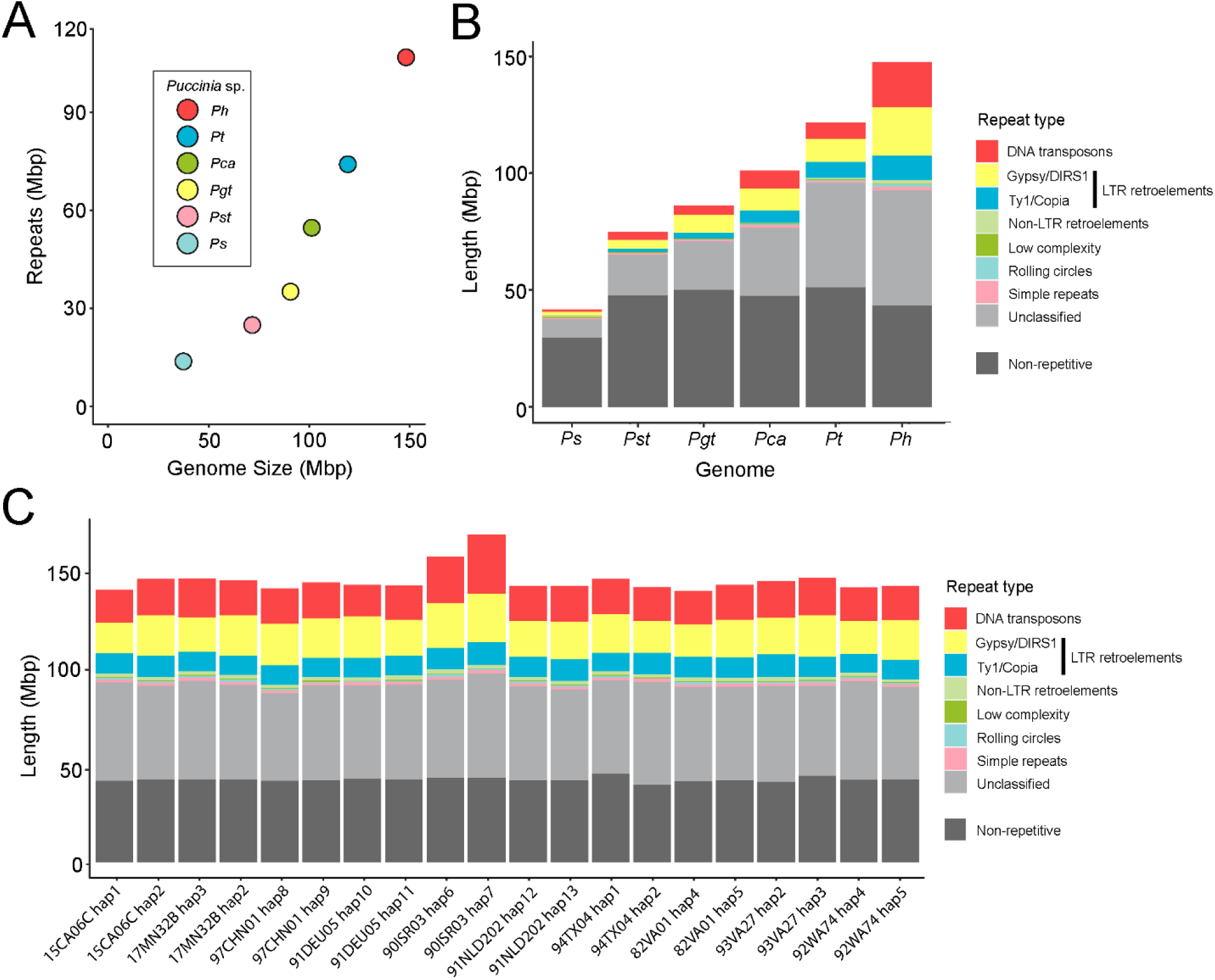
A) Genome sizes (Mbp) of a single representative chromosome-level nuclear haplotype from *Puccinia* species (*Ps*= *P. silphi* haplotype 1 (de Mattos et al., 2025), *Pst* = *P. striiformis* f. sp. *tritici* AZ2 hap1 (Wang et al., 2024), *Pgt* = *P. graminis* f. sp. *tritici* ETH2013-1 hap4 (Spanner et al. 2025), *Pca* = *P. coronata* f. sp. *avenae* 203 hapA (Henningsen et al., 2024), *Pt* = *P. triticina* 19NSW04 hapB (Sperschneider et al., 2023), *Ph* = *P. hordei* 17MN32B hap2), and length of genome in Mbp covered by repeat sequences. B) Length of genome in Mbp covered by distinct types of repeat element in haplotype-phased genome assemblies from the different *Puccinia* species presented in (A). C) The length of the haploid nuclear-phased *P. hordei* genomes covered by distinct types of repeat element for core chromosomes 1-18.

### Establishment of a *P. hordei* pan-genome and pan-effectorome

A range of 36,203 to 37,610 protein-coding genes were annotated in the diploid *P. hordei* genomes (**Table S23**). An average of 17,841 genes were annotated per nuclear haplotype. This was higher than the average of 11,309 reported by Yu et al. (2025) for phased *P. hordei* genomes but comparable to the number of genes annotated in fully phased genomes of *P. graminis* f. sp. *tritici* (Li et al., 2019), *P. triticina* (Sperschneider et al., 2023) and *P. coronata* f. sp. *avenae* (Henningsen et al., 2024). The mean BUSCO v3 completeness score for haploid gene annotations was 90.3% with a mean combined haplotype score of 93.9% (**Table S24**). Approximately 21% of the genome was covered in gene-coding regions and the mean length of genes was ∼1.7 kb (**Table S23**).

A unique set of 13 nuclear haplotypes was analyzed as the global *P. hordei* pan-genome, including 94TX04 hap1, 93VA27 hap2, 93VA27 hap3, 82VA01 hap4 and 82VA01 hap5 as representative of haplotypes that are shared between multiple isolates. To identify core and non-core genes in the *P. hordei* pangenome, we conducted an orthogroup analysis on the proteome derived from the longest isoform of all gene annotations in these haplotypes. The Heaps’ Law decay parameter for pan-genome growth (α) was greater than 1 (α =1.679), indicating a closed pan-genome with minimal new gene family discovery on addition of novel nuclear haplotypes. This can be visualized as a plateauing saturation curve in **Figure 6A**. The Chao lower bound estimated a minimum of 19,888 total protein orthogroups in the pan-genome, suggesting that the current haplotype collection may have captured up to 99.96% of protein orthogroups. Of 19,881 total protein orthogroups: 7,092 (35.7%) were shared by all 13 haplotypes (core), 2,834 (14.3%) were shared by 11-12 haplotypes (soft core), 6,248 (31.4%) were shared by 4-10 haplotypes (shell), 3,533 (17.8%) were shared by 2-3 haplotypes (cloud) and only 174 (0.9%) were haplotype-specific (**Figure 6B**).

**Figure 6.**
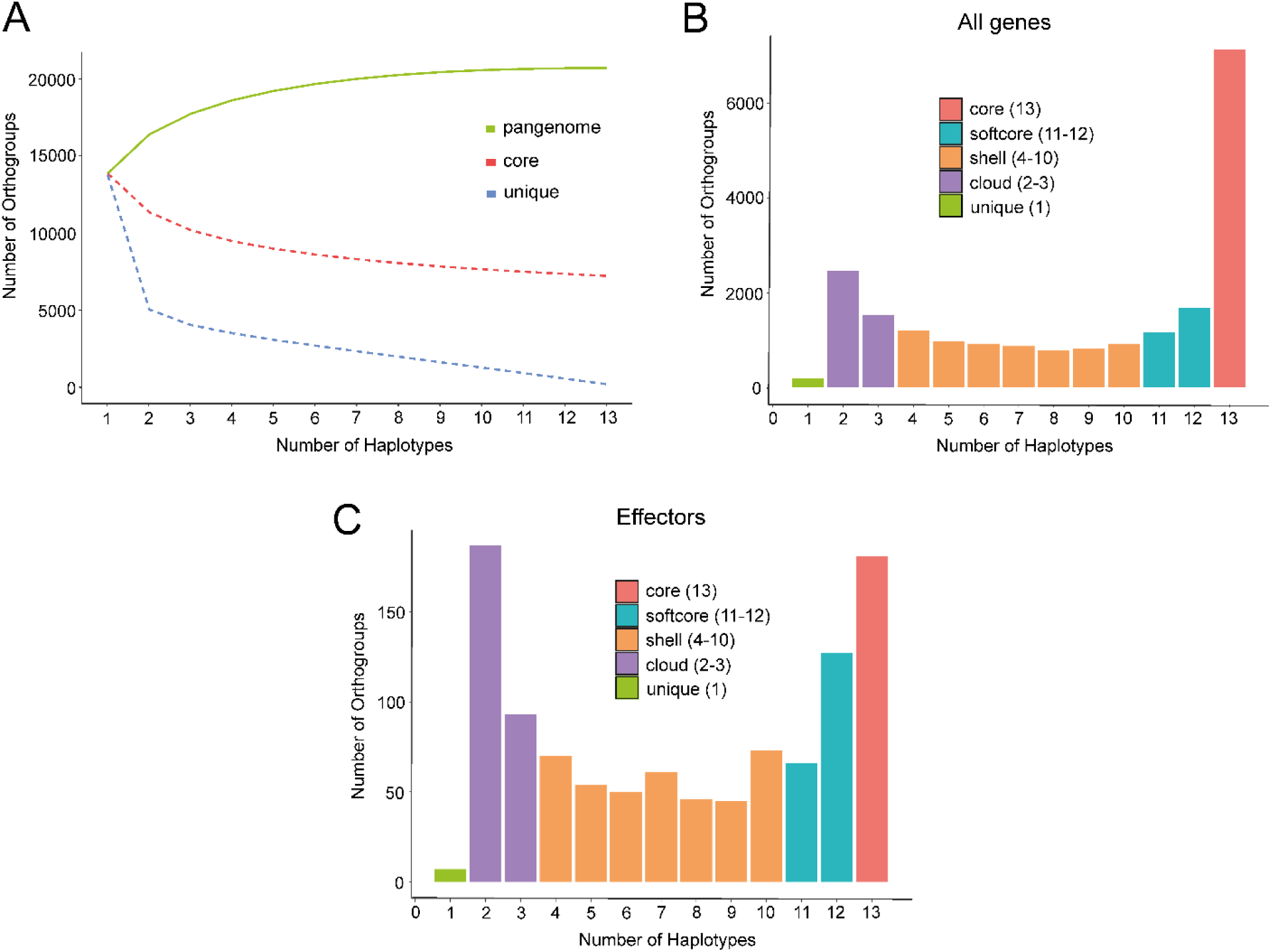
A) Rarefaction analysis of the *P. hordei* pan-genome based on orthogroup presence/absence across the 13 unique nuclear haplotypes. The number of core, unique, and total (pan-genome) orthogroups was calculated for 1000 random permutations of haplotype order at each sampling depth from 1 to 13 haplotypes. Lines show the mean number of orthogroups across permutations per category for each haplotype value. B) Number of gene orthogroups classified as core (shared by all 13 nuclear haplotypes), softcore (shared by 11-12 haplotypes), shell (shared by 4-10 haplotypes), cloud (shared by 2-3 haplotypes) and unique (haplotype-specific). C) Number of predicted effector orthogroups classified as core (shared by all 13 nuclear haplotypes), softcore (shared by 11-12 haplotypes), shell (shared by 4-10 haplotypes), cloud (shared by 2-3 haplotypes) and unique (haplotype-specific).

Each haplotype encoded 1,566-1,714 predicted secreted proteins with 629-731 predicted effectors. The predicted effectors from the same unique set of 13 nuclear haplotypes were analyzed as the pan-effectorome. A total of 973 effector orthogroups were identified. Of these, 175 (18.0%) orthogroups were shared between all haplotypes (core), 199 (20.5%) orthogroups were shared between 11-12 haplotypes (soft core), 367 (37.7%) orthogroups were shared between 4-10 haplotypes (shell), 226 (23.2%) orthogroups were shared between 2-3 haplotypes and there were 6 (0.6%) haplotype-specific orthogroups (**Figure 6C**).

## Discussion

Pan-genomic analyses offer a powerful framework for exploring evolutionary mechanisms and genetic diversity underpinning pathogen populations on a global scale. This has been particularly important for dikaryotic species in the genus *Puccinia*, where recent studies have revealed that entire nuclei can be exchanged between isolates generating novel genetic diversity (Henningsen et al., 2024; Li et al., 2019; Sperschneider et al., 2023). Through genomics-based surveillance, we can track the movement of entire nuclei in global *Puccinia* populations. In this study, we developed a set of ten high-quality annotated haplotype-phased genome assemblies for *P. hordei* capturing a sample of genetic diversity in the US as well as several other countries over a period of 35 years. Comparative genomics based on this *Ph* pan-genome revealed that multiple somatic hybridization events had likely occurred to generate the observed global lineages. Notably, hap2 was shared by two distinct clonal lineages in the US and was also contained in an Australian lineage (isolate Ph560) (Chen et al., 2019). It is also possible that Ph560 is related to hap2 by a single meiotic recombination event in which the two nuclei of Ph560 as the sexual parent recombined to give rise to hap2, as has been observed for a field derived F_1_ progeny isolate in *P. coronata* f. sp. *avenae* (Henningsen et al., 2024). Comparison of hap2 with the fully phased genome of Ph560 (Yu et al., 2025) would be required to confirm nuclear haplotype identity. These results demonstrate the importance of clonal propagation and nuclear exchange in global population dynamics of *P. hordei*.

The lack of the sexual reproduction cycle of *P. hordei* in the US is apparent with the presence and long-term persistence of multiple clonal lineages e.g. lineage 1 from 1994-2015; lineage 2 from 1993-2017; lineage 3 from 1982-1992. This was expected given the scarcity of the main alternate host (*Ornithogalum* spp.) near barley production areas. Even where the alternate host is present in parts of the country (e.g. Virginia), the timing of *P. hordei* teliospore formation may not coincide with the growth cycle of *Ornithogalum*, restricting opportunities for sexual recombination (Clifford, 1985). Consistent with the tetrapolar mating system observed in other *Puccinia* species (Henningsen et al., 2024; Luo et al., 2024; Sperschneider et al., 2023), all isolates exhibited divergent alleles between dikaryotic nuclei at the two mating type loci *PR* (*a*) and *HD* (*b, bWbE*). The *PR* locus appeared bi-allelic, whilst the *HD* locus was multiallelic. Analyses of the homeodomain transcription factor locus (*bWbE*) demonstrated that isolates sharing nuclear haplotypes also shared *bW* and *bE* alleles (e.g. *bW2* and *bE2* for isolates sharing HapB). We also found that hap1, hap4, hap10 and hap12 (possessed by isolates from US, Netherlands and Germany) shared the *bW1* and *bE1* alleles. It is possible that these haplotypes had a common ancestral nuclear haplotype which underwent recombination via meiosis to generate these distinct nuclear haplotypes. This demonstrates that sexual recombination is another important, although seemingly rare, evolutionary mechanism generating novel diversity in *P. hordei*.

We also found that mitochondria were identical in all ten isolates, further supporting long-term clonal propagation across global populations. The uniparental inheritance (or fixation of mitochondrial genome) in this group of isolates suggests that all are descendants of a single maternal lineage that has expanded due to agriculture. International migration of *P. hordei* lineages must have occurred leading to the presence of isolates sharing nuclei on distinct continents e.g. isolates from the USA and Australia share hap2. Exotic incursions of multiple distinct foreign *Puccinia* isolates have been reported previously (e.g. *P. triticina* in Australia (Sperschneider et al., 2023) and *P. graminis* f. sp. *tritici* in Ethiopia (Olivera et al., 2015)), and while airborne dispersal of urediniospores is a common mode of spread for these pathogens, people who travel internationally with contaminated clothing can also introduce new pathogen strains into different countries (Wellings 2007).

Rust fungi are renowned for their large genomes caused by expansion of repetitive elements occupying ∼45-90% of the genome (Corre et al., 2025; Gupta et al., 2023; Lorrain et al., 2019). We found that the increase in size of barley leaf rust genomes when compared to other cereal rust pathogens could be explained by a larger proportion of the genome covered by repetitive elements (∼70%), but with a similar sized non-repetitive genome component. Similarly to Corre et al. (2025), we found that the largest class of annotated TEs were Class I (retrotransposons), followed by Class II (DNA transposons). Also in agreement with Corre et al. (2025), Long-Terminal Repeat (LTR) retrotransposons were the most abundant and of these, Gypsy/TIR1 dominated. Any intraspecific variation in these classes (e.g. 90ISR03) may represent recent bursts of transposition. Accumulation of TEs is an ancestral trait in rust fungi, which may have evolved to tolerate this high TE load by mechanisms distinct from Repeat-Induced Point (RIP) mutations which are present in the ascomycete fungi (Cambareri et al., 1991; Corre et al., 2025; Clutterbuck, 2011). Since the DNA methyltransferase genes DNMT1 and DNMT5 are present, it is plausible that TE silencing could occur via DNA methylation.

We observed three distinct scenarios of reciprocal interchromosomal translocations within the nuclear haplotypes. It is well-known that repeat elements can facilitate such chromosomal rearrangements, and that they occur more frequently during meiosis than mitosis (Jinks-Robertson & Petes, 1986). While the precise mechanism of rearrangement within these *P. hordei* isolates remains unclear, the presence of distinct chromosomal configurations among isolates with otherwise identical haplotypes suggests that at least two of the three translocations arose during clonal propagation rather than through meiotic recombination. Fouché et al. (2023) and Badet et al. (2024) pinpointed a single transposable element, DNA transposon *Styx*, as the trigger for chromosome instability and degeneration in *Zymoseptoria tritici*. It is possible that specific transposable elements are responsible for the chromosomal rearrangements observed in *P. hordei*, but further studies are required to improve repeat element annotation. Translocation between two chromosomes within a nuclear haplotype has also been observed in both *P. graminis* f. sp. *tritici* and *P. triticina* (Li et al., 2019; Sperschneider et al., 2023), suggesting that such structural variation may be common in highly clonal species of *Puccinia*.

The distinct genome architecture of isolate 90ISR03 (genome size, repeat content, chromosome number) and sequence divergence from the other isolates raises interesting taxonomic questions regarding *Puccinia hordei*. In *Puccinia* species, distinct *formae speciales* (Anikster, 1984b) are distinguished by their host specialization-sub-species that can infect multiple plant species, often with overlapping host ranges (Huang et al., 2019). Like the other *P. hordei* isolates, isolate 90ISR03 infects and completes its asexual cycle (urediniospores) on cultivated and wild barley (*Hordeum vulgare* subsp. *spontaneum*). Phenotypically, isolate 90ISR03 can only be distinguished from other *P. hordei* isolates by its ability to overcome resistance gene *Rph15*. However, we do not know if all isolates are able to complete their sexual cycle on species of *Ornithogalum*, *Leopoldia* or *Dipcadi*. It has previously been shown that both wild and cultivated grasses were important in the evolution of new strains of *P. graminis*, suggesting the formation of somatic hybrids between *P. graminis* f. sp. *tritici* and *P. graminis* f. sp. *secalis* (the rye stem rust pathogen) (Luig & Watson, 1972). Isolate 90ISR03 could represent a distinct *formae speciales* to the other isolates tested, but more infection tests are required to clearly define this (e.g. establish uredinial host range and determine if it can complete the sexual cycle on the same alternate host). It could also be a cryptic species where it is indistinguishable from other *P. hordei* phenotypically, but genetically distinct. 90ISR03 was sampled in Israel, where populations of *H. vulgare* subsp. *spontaneum* and other wild *Hordeum* species are common. There is more potential for novel lineages to evolve, whilst hybridizing with distinct *formae speciales* and/or specializing to infect distinct sets of host species. Sequencing and phenotyping of more *P. hordei* isolates from this region would give a clearer picture of the genetic diversity present.

Long-read sequencing permits us to analyze the telomere-to-telomere chromosome-level structure of genomes, which in turn can reveal structural abnormalities. The 90ISR03 assembly harbored an additional scaffold (designated chr19) in hap6 containing hallmarks of a chromosome with a centromeric repeat region and two telomeres. Accessory chromosomes (or “supernumerary” or “lineage-specific” chromosomes) have been well described in ascomycete fungi (Mehrabi et al., 2011). These chromosomes are typically smaller than core chromosomes, may have a higher repeat density and are not essential for the survival of the organism (Habig & Stukenbrock, 2020). However, in plant pathogenic fungi, they have often been shown to give a selective advantage by harboring host-specific toxin genes (Harimoto et al., 2007; Witte et al., 2021), detoxification genes (Han et al., 2001) or small secreted effector genes (Ayukawa et al., 2021; Peng et al., 2019). In the case of isolate 90ISR03, the extra scaffold is comparable in size to the core chromosomes, is repeat-rich, and harbors effector-like genes. Sequencing of additional isolates from the same host cultivar and region may help to clarify how it was acquired and its biological role. Henningsen et al. (2025) also identified a supernumerary chromosome in a single oat crown rust (*P. coronata* f. sp. *avenae*) isolate from Israel. Similarly to chr19 in *P. hordei* 90ISR03, the scaffold was present in a single nuclear haplotype, had comparable HiFi read depth coverage to core chromosomes and encoded putative effector genes. In contrast, the oat crown rust scaffold was smaller than core chromosomes (2.17Mbp) and had comparable repeat content to core chromosomes. Together, these observations suggest that accessory chromosomes may be a common feature among *Puccinia* species.

To further explore genome content variation within the species, we curated a pan-genome for *P. hordei*. Heaps’ Law analysis suggested that the pan-genome comprised of 13 unique *P. hordei* haplotypes had approached saturation when assessing the accumulation of novel gene families. We further annotated the putative effectorome using a combination of infection time point RNAseq data and bioinformatics prediction. Recognized effectors (avirulence proteins) are yet to be identified in *P. hordei*. These phased annotated genomes can be utilized as reference sequences for the identification of novel avirulence genes via the sequencing of diverse isolates (association mapping) or mutation screening of clonal lineages.

### Conclusions

This study presents haplotype-phased genome assemblies for *P. hordei*, providing critical insights into the global population dynamics, genome structure, and evolutionary mechanisms of this important barley pathogen. Our pan-genomic approach uncovered evidence for widespread clonal propagation, and nuclear exchange shaping *P. hordei* diversity, particularly in regions lacking the alternate host required for sexual reproduction. Conserved mitochondrial genomes and shared mating type alleles further support rare recombination events and long-term clonal expansion of a few founding lineages. Structural genome variation, including chromosomal translocations and a potential accessory chromosome, highlights the plasticity and adaptability of *P. hordei* genomes, likely mediated by transposable elements. The distinct genomic architecture of isolate 90ISR03 raises questions about *formae speciales* boundaries and suggests a need for further taxonomic investigation. Together, this set of annotated phased genomes represent a foundational resource for future studies on the evolution of global *P. hordei* lineages and will support ongoing efforts in pathogen surveillance.

## Declarations

### Ethics approval and consent to participate

Not applicable.

### Consent for publication

Not applicable.

### Data availability

Raw reads generated in this study are available at NCBI under BioProject PRJNA1298276. All haplotype-phased assemblies and gene annotations are available at doi: 10.6084/m9.figshare.30623441. Code used for data visualization can be found in the github repository: https://github.com/bspan07/puccinia_hordei_pangenome.

## Competing interests

The authors declare that they have no competing interests.

## Funding

This project was partially supported by USDA-NIFA BBSRC award 2022-67013-36505 and the Lieberman-Okinow Endowment at the University of Minnesota.

## Authors’ contributions

**Rebecca Spanner:** Data curation, Formal analysis, Investigation, Methodology, Project administration, Software, Visualization, Writing – original draft, Writing – review & editing

**Eric Nazareno:** Data curation, Formal analysis, Investigation, Writing - original draft, Writing - review & editing

**Eva C. Henningsen:** Investigation, Methodology, Writing - review & editing

**Alexis Feist:** Investigation, Writing - review & editing

**Matthew J. Moscou:** Funding acquisition, Supervision, Writing - review & editing

**Matthew N. Rouse:** Funding acquisition, Resources, Supervision, Writing - review & editing

**Hayley Mangelson**: Formal analysis, Writing – review & editing

**Kyle Langford:** Investigation, Writing – review & editing

**Ivan Liachko:** Funding acquisition, Methodology, Resources, Supervision, Writing – review & editing

**Jibril Lubega:** Investigation, Resources, Writing - review & editing

**Kostya Kanyuka:** Conceptualization, Funding acquisition, Resources, Supervision, Writing - review & editing

**Jana Sperschneider:** Methodology, Software, Supervision, Writing - review & editing

**Peter N. Dodds:** Conceptualization, Funding acquisition, Project administration, Resources, Supervision, Writing - review & editing

**Melania Figueroa:** Conceptualization, Funding acquisition, Project administration, Resources, Supervision, Writing - review & editing

**Brian J. Steffenson:** Conceptualization, Funding acquisition, Project administration, Resources, Supervision, Writing - review & editing

## Acknowledgements

We thank Emmery Hartwig for technical assistance with the increase of barley leaf rust isolates; Stephanie Dahl and Aubree Kees for their assistance at the University of Minnesota Biosafety Level 3 containment facility; and the Minnesota Supercomputing Institute for computing resources.

## Notes

https://doi.org/10.6084/m9.figshare.30623441

https://www.ncbi.nlm.nih.gov/bioproject/PRJNA1298276/

### Competing Interest Statement

The authors have declared no competing interest.

