## Supplementary Figures for "Haplotype-phased genomes of the barley leaf rust pathogen reveal evidence of repeat element expansion and somatic hybridization"

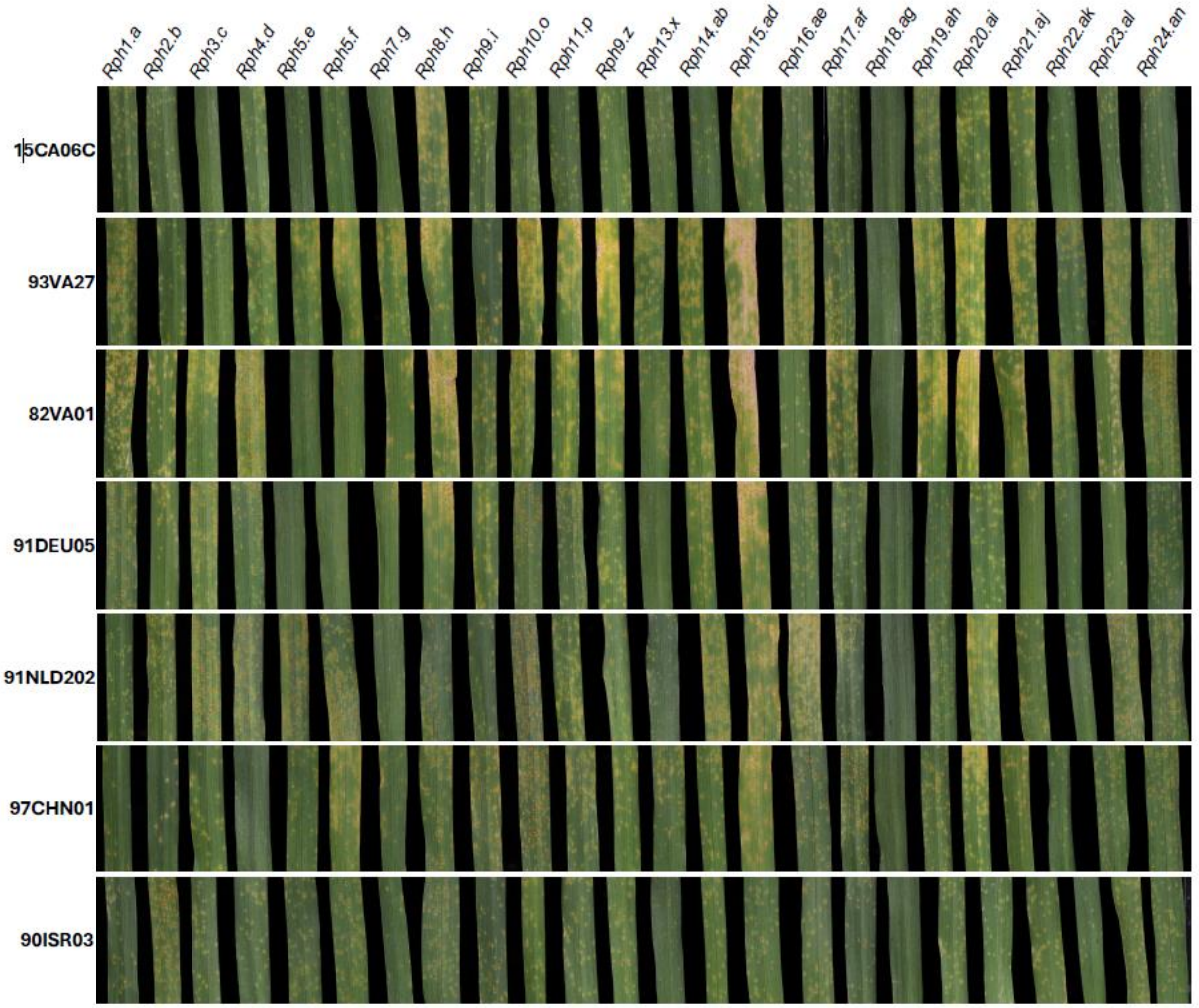


**Figure S1.** Phenotypes of representative *Puccinia hordei* isolates from each of the seven lineages on the *Rph1* to *Rph24* barley leaf rust differential set.

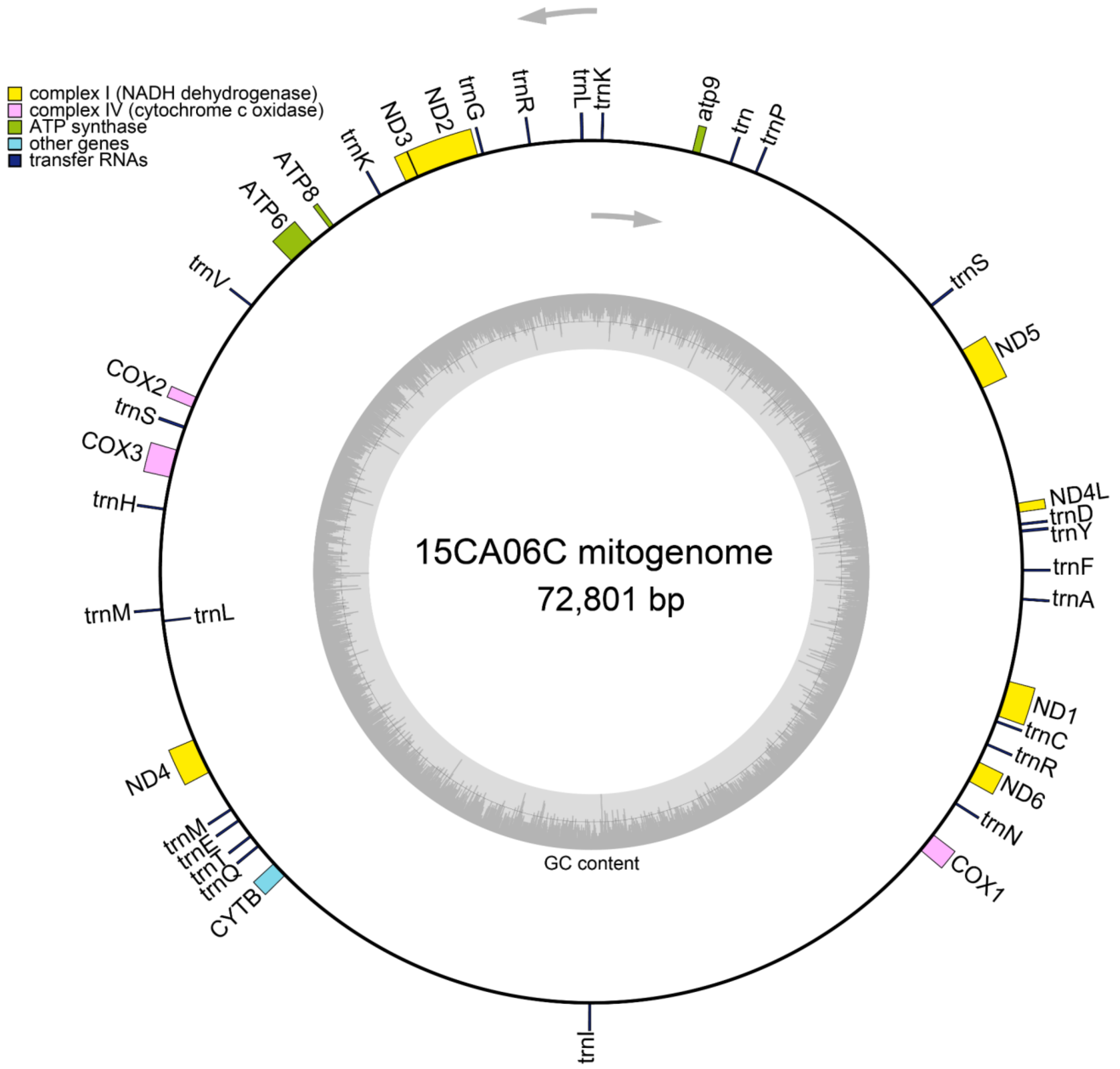


**Figure S2.** Genome map of the mitochondrial assembly for *Puccinia hordei* isolate 15CA06C (rotated PacBio HiFi contig ptg000045l) annotated with 39 mitochondrial genes (color-coded by function). The direction of transcription is denoted by the arrows. GC content of mitochondrial sequence is displayed as an inner circular plot.

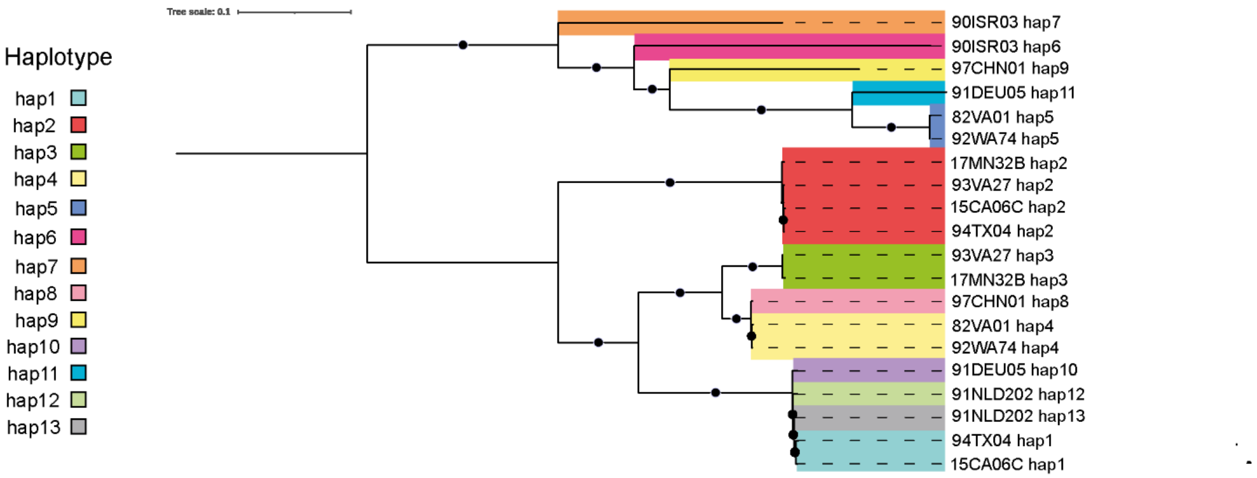


**Figure S3.** Maximum likelihood tree with 500 bootstraps showing the phylogenetic relationship between the 20 *P. hordei* haploid genomes, color-coded by nuclear haplotype identity (hap1-hap13), after alignment to isolate 15CA06C hap2. Branches with circular node are supported by >80% of bootstraps.

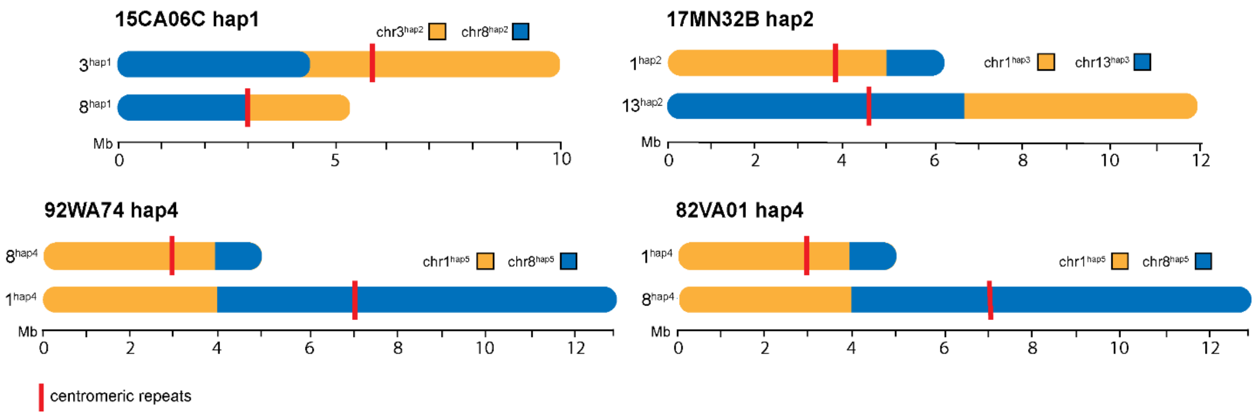


**Figure S4.** Schematic of the structurally distinct chromosomes in the collection of haplotype-phased *P. hordei* assemblies. Chromosomes are color-coded by their sequence homology to the corresponding chromosomes in the other haplotype within that isolate. The approximate position of alpha-satellite repeats is marked in red and was inferred through inspection of chromatin contact maps.

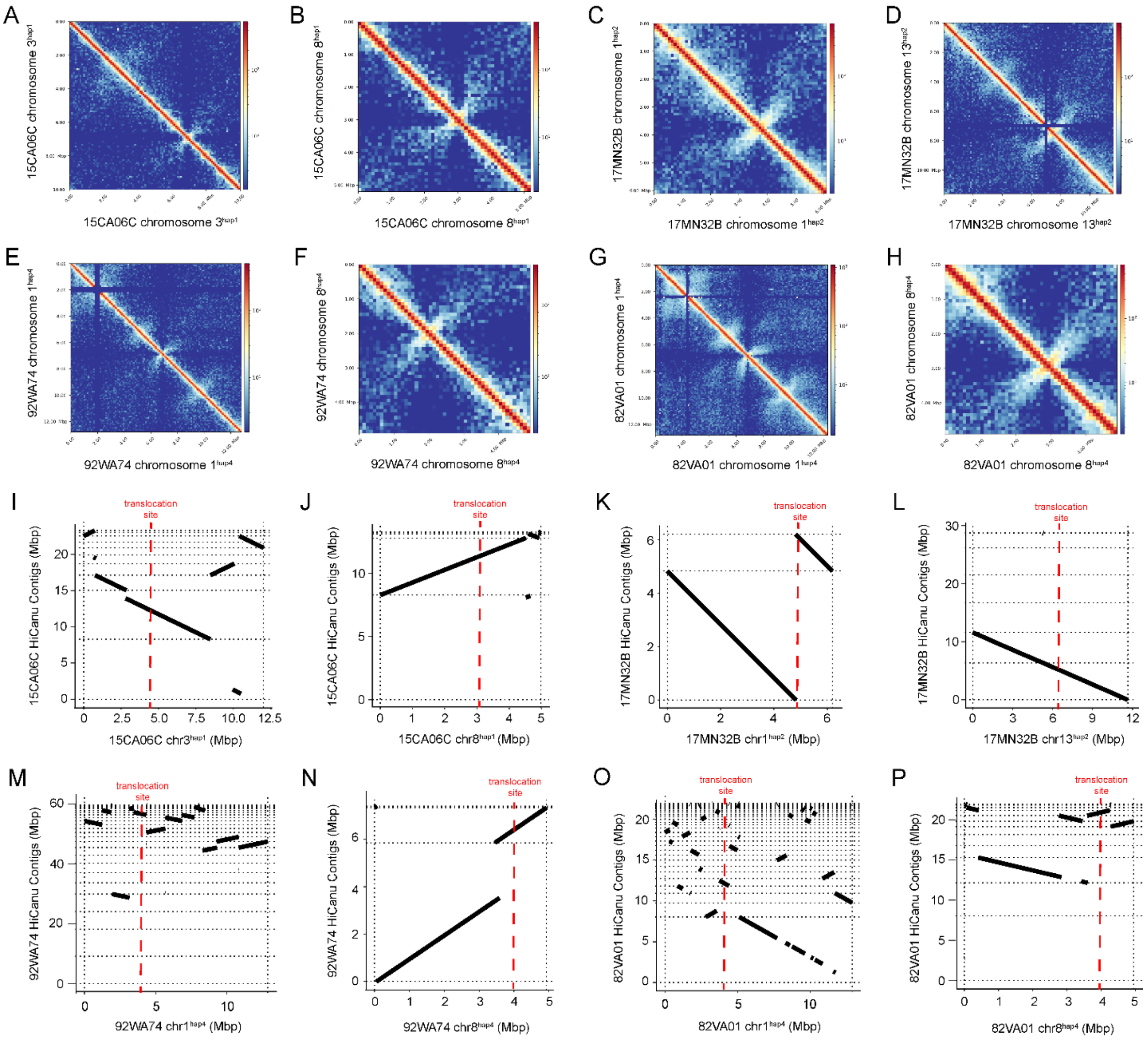


**Figure S5**. A-H) Hi-C chromatin contact maps of *P. hordei* chromosomes with translocations. I-P) Chromosomes with translocations were aligned to HiCanu contigs for the corresponding isolate using D-Genies.

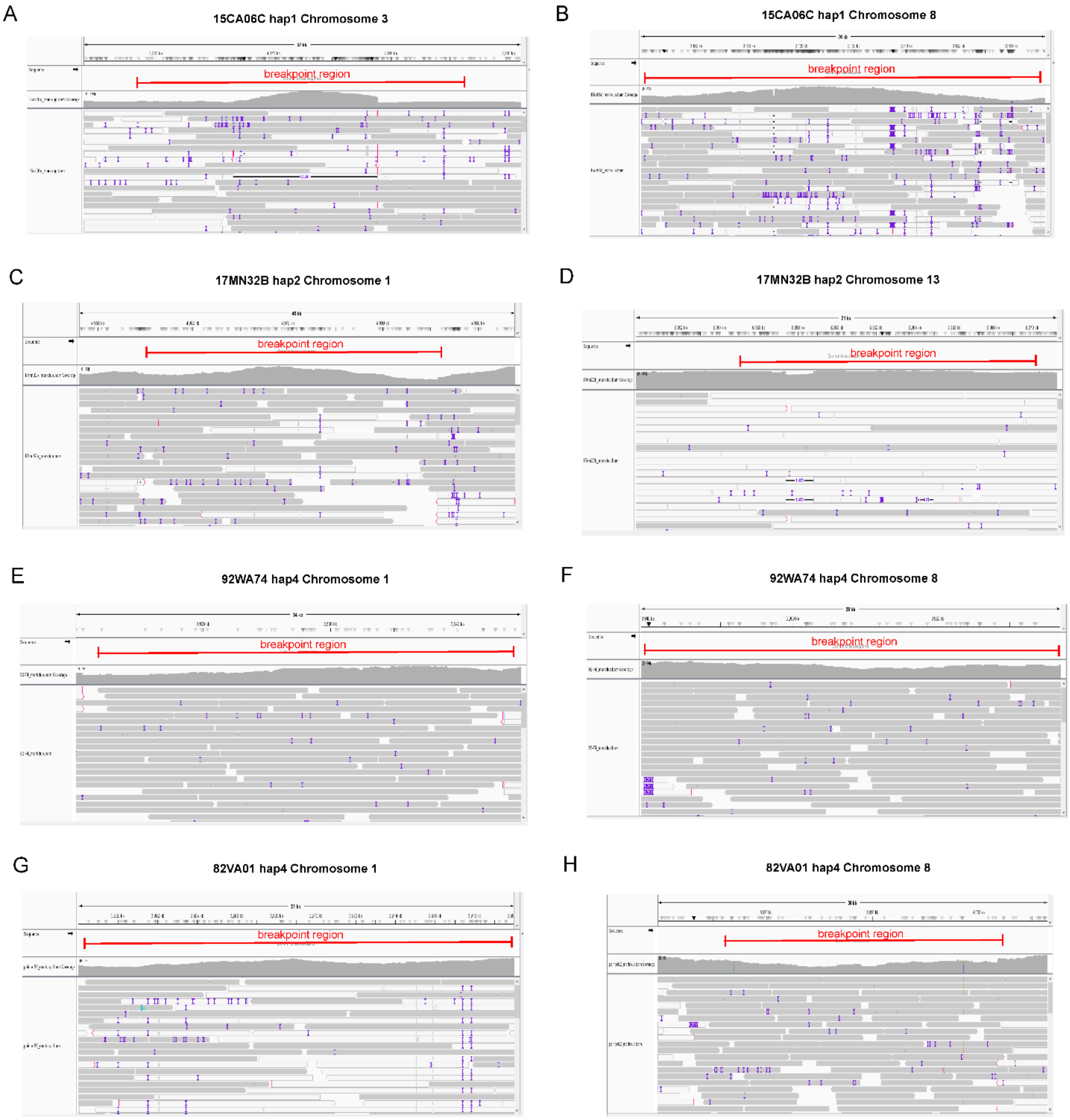


**Figure S6.** Raw PacBio HiFi reads mapping (no multi-mapping) across breakpoint regions determined from D-Genies pairwise alignments of homologous chromosomes (translocated vs. non-translocated) in *Puccinia hordei*.

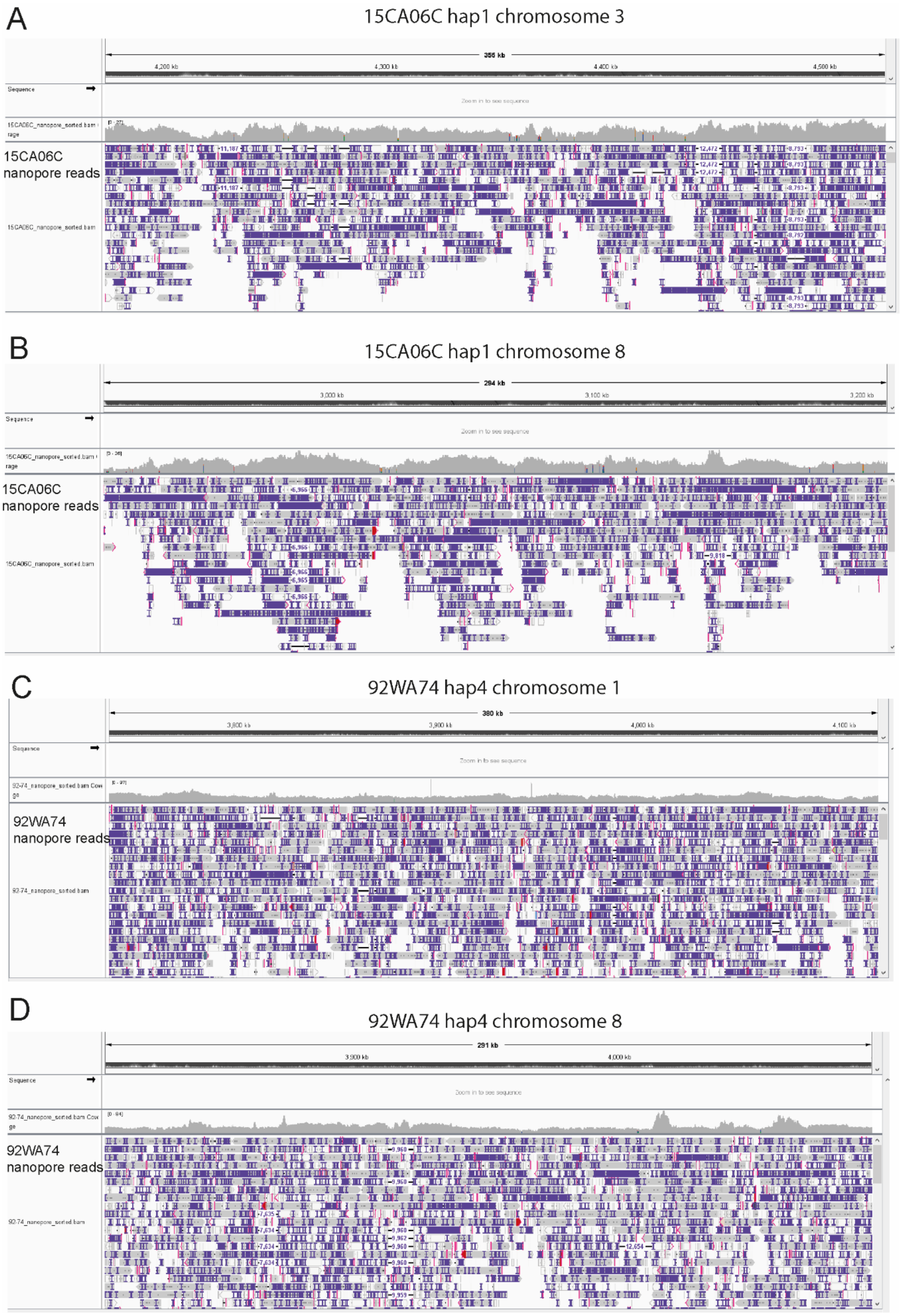
A
15CA06C hap1 chromosome 3
356 kb
4,200 kb
4,300 kb
4,400 kb
4,500 kb
Sequence
Zoom in to see sequence
15CA06C nanopore reads
B
15CA06C hap1 chromosome 8
294 kb
3,000 kb
3,100 kb
3,200 kb
Sequence
Zoom in to see sequence
15CA06C nanopore reads
C
92WA74 hap4 chromosome 1
380 kb
3,800 kb
3,900 kb
4,000 kb
4,100 kb
Sequence
Zoom in to see sequence
92WA74 nanopore reads
D
92WA74 hap4 chromosome 8
291 kb
3,900 kb
4,000 kb
Sequence
Zoom in to see sequence
92WA74 nanopore reads

**Figure S7.** Nanopore reads aligned (no multi-mapping) to phased diploid assembly with the approximate translocation breakpoint visualized using Integrated Genome Viewer for A) isolate 15CA06C hap1 chromosome 3, B) isolate 15CA06C hap1 chromosome 8, C) isolate 92WA74 hap4 chromosome 1 and D) isolate 92WA74 hap4 chromosome 8 of *Puccinia hordei*.

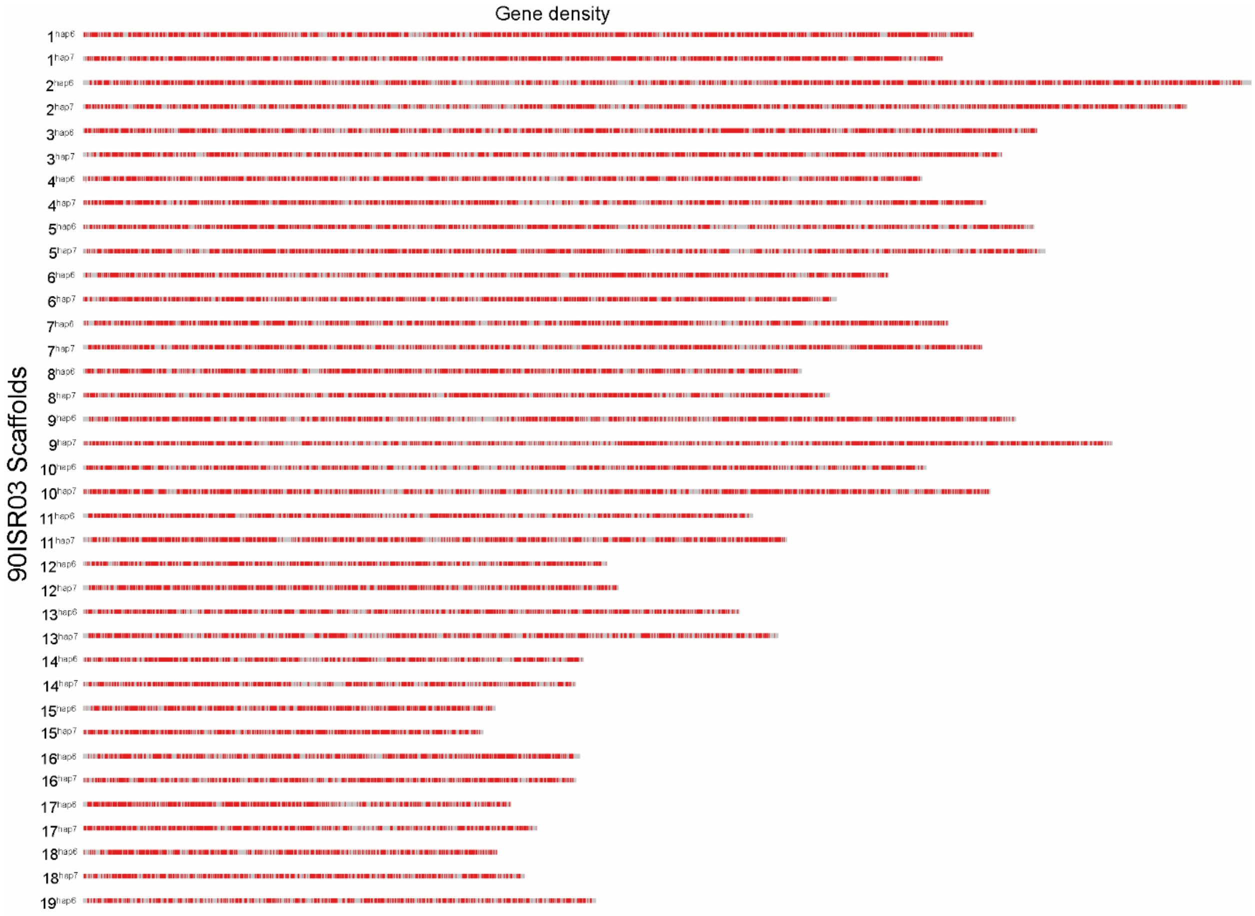


**Figure S8**. A) Chromosome-level map showing gene density of the chromosomes in each of the phased nuclear haplotypes, hap6 (19 scaffolds) and hap7 (18 scaffolds), of *P. hordei* isolate 90ISR03.

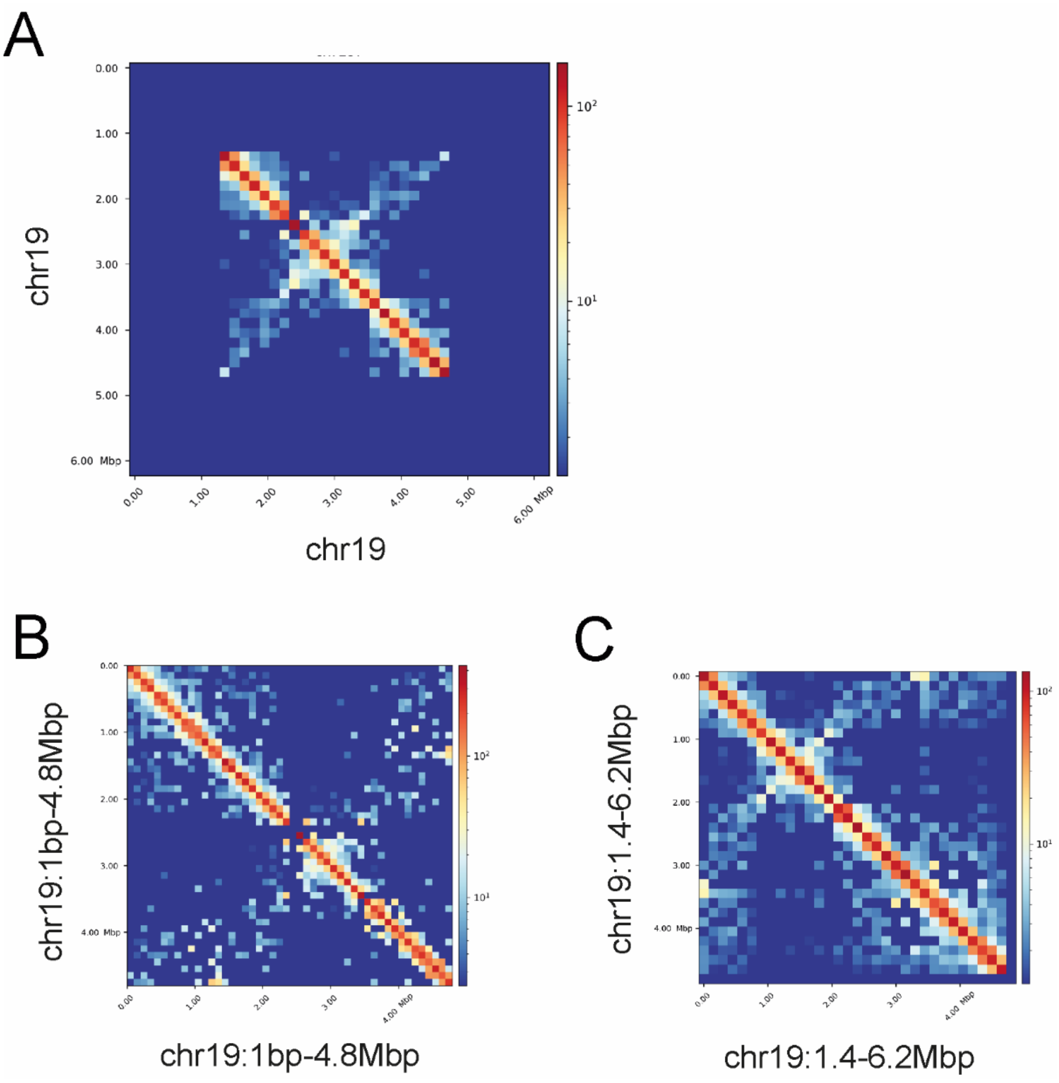


**Figure S9.** Chromosome-level HiC chromatin contact maps for A) full-length 6.2Mbp chromosome 19, B) the first 4.8Mbp of chromosome 19 and C) the last 4.8Mbp (1.4-6.2Mbp) of chromosome 19, in hap6 of *P. hordei* isolate 90ISR03.

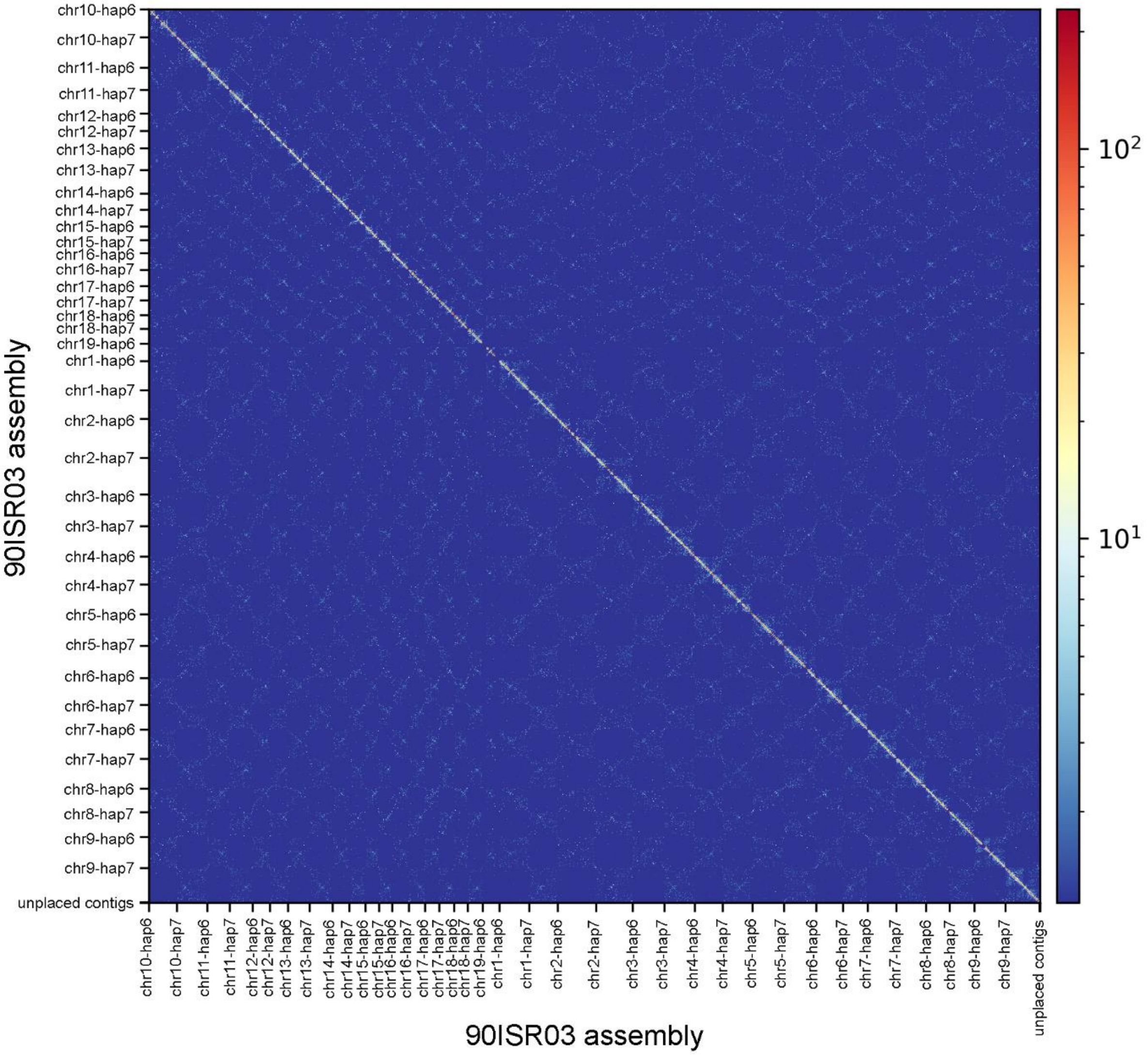


**Figure S10.** HiC chromatin contact map for the diploid 90ISR03 assembly.

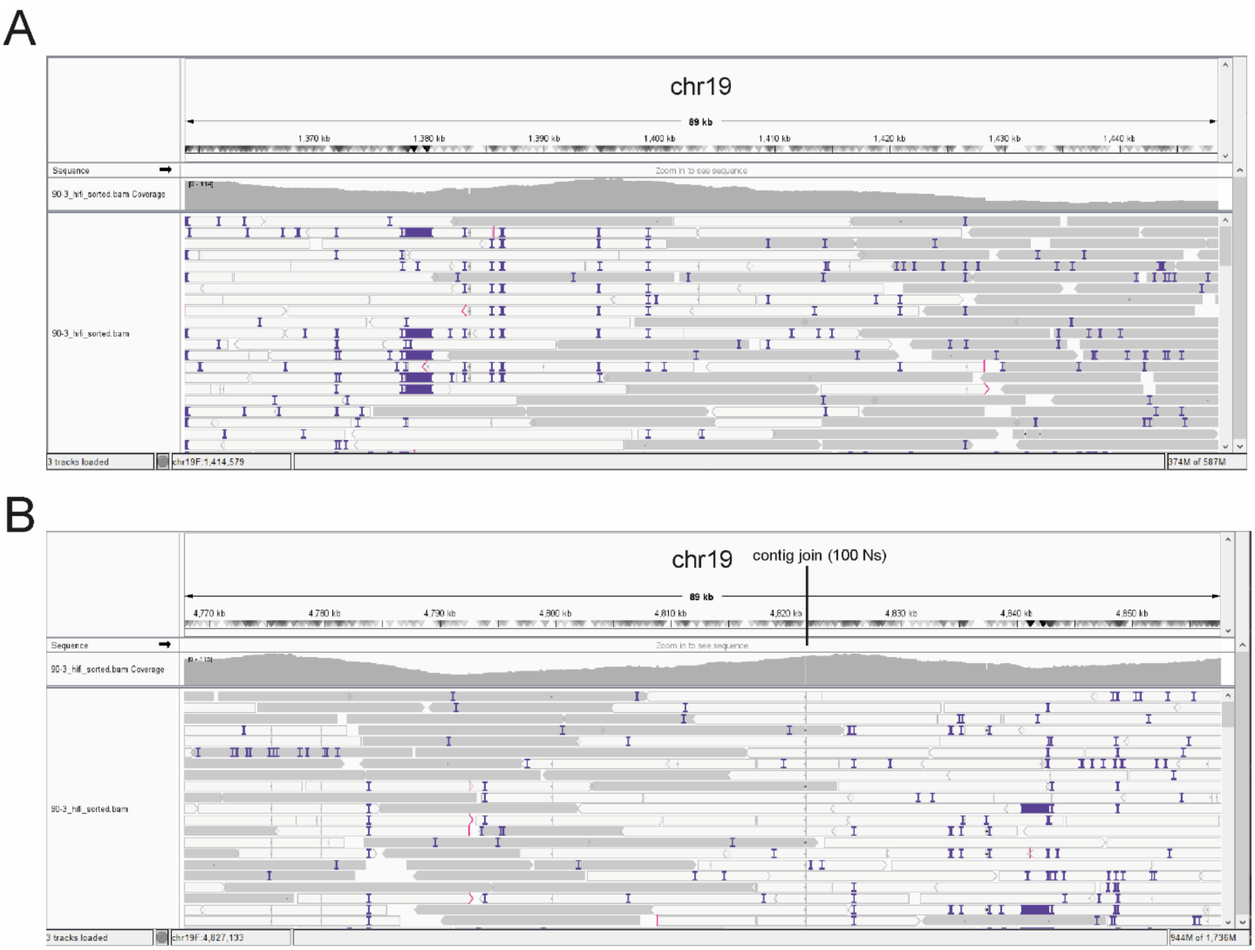


**Figure S11.** PacBio HiFi reads mapping to chr19 in hap6 of 90ISR03 A) over the boundary of the first duplicated ~1.4Mbp region, and B) over the boundary where the two constituent contigs of chr19 are joined (100 N’s) and where the second duplicated ~1.4Mbp region begins.

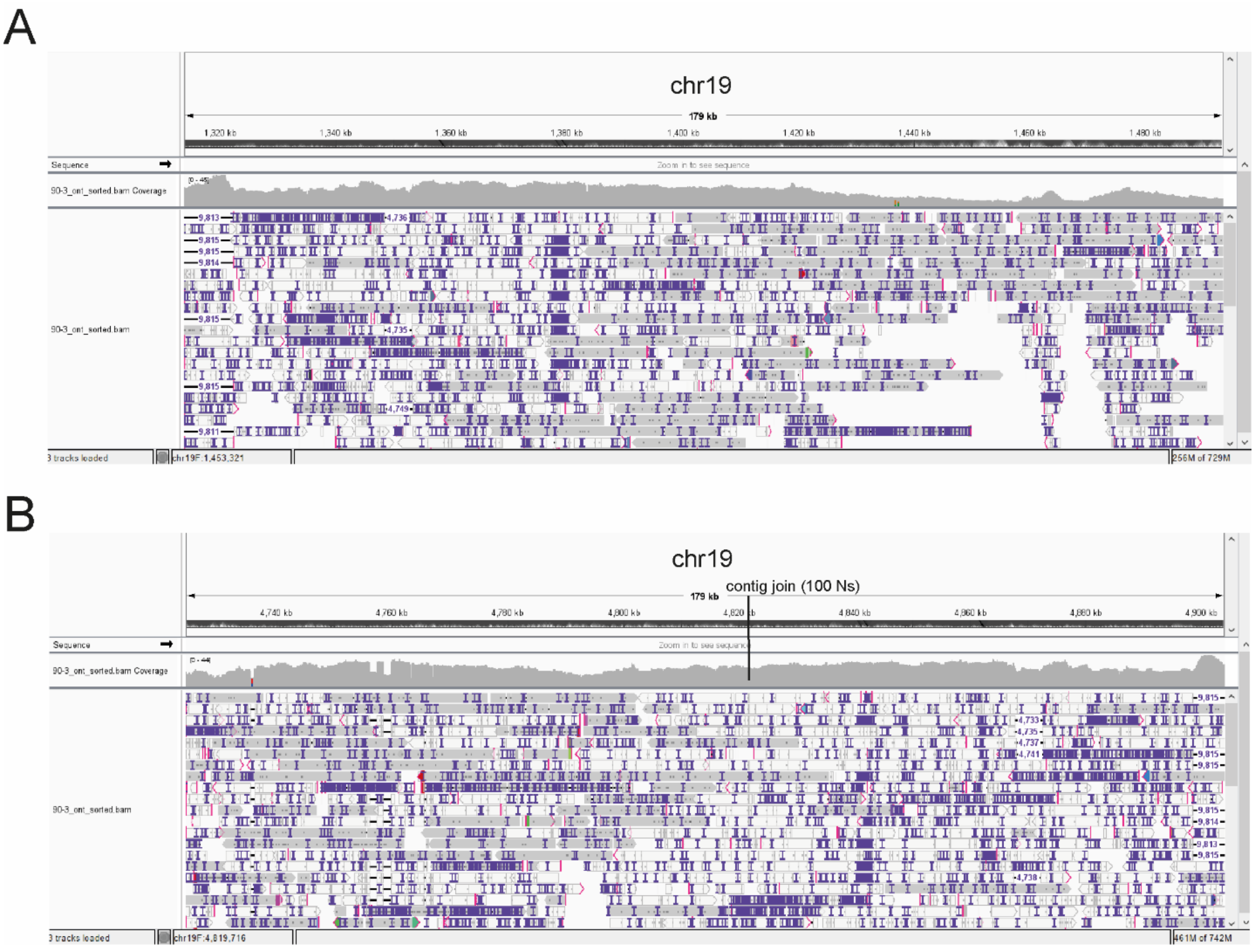


**Figure S12.** Oxford nanopore technology reads mapping to chr19 in hap6 of 90ISR03 A) over the boundary of the first duplicated ~1.4Mbp region, and B) over the boundary where the two constituent contigs of chr19 are joined (100 N's) and where the second duplicated ~1.4Mbp region begins.

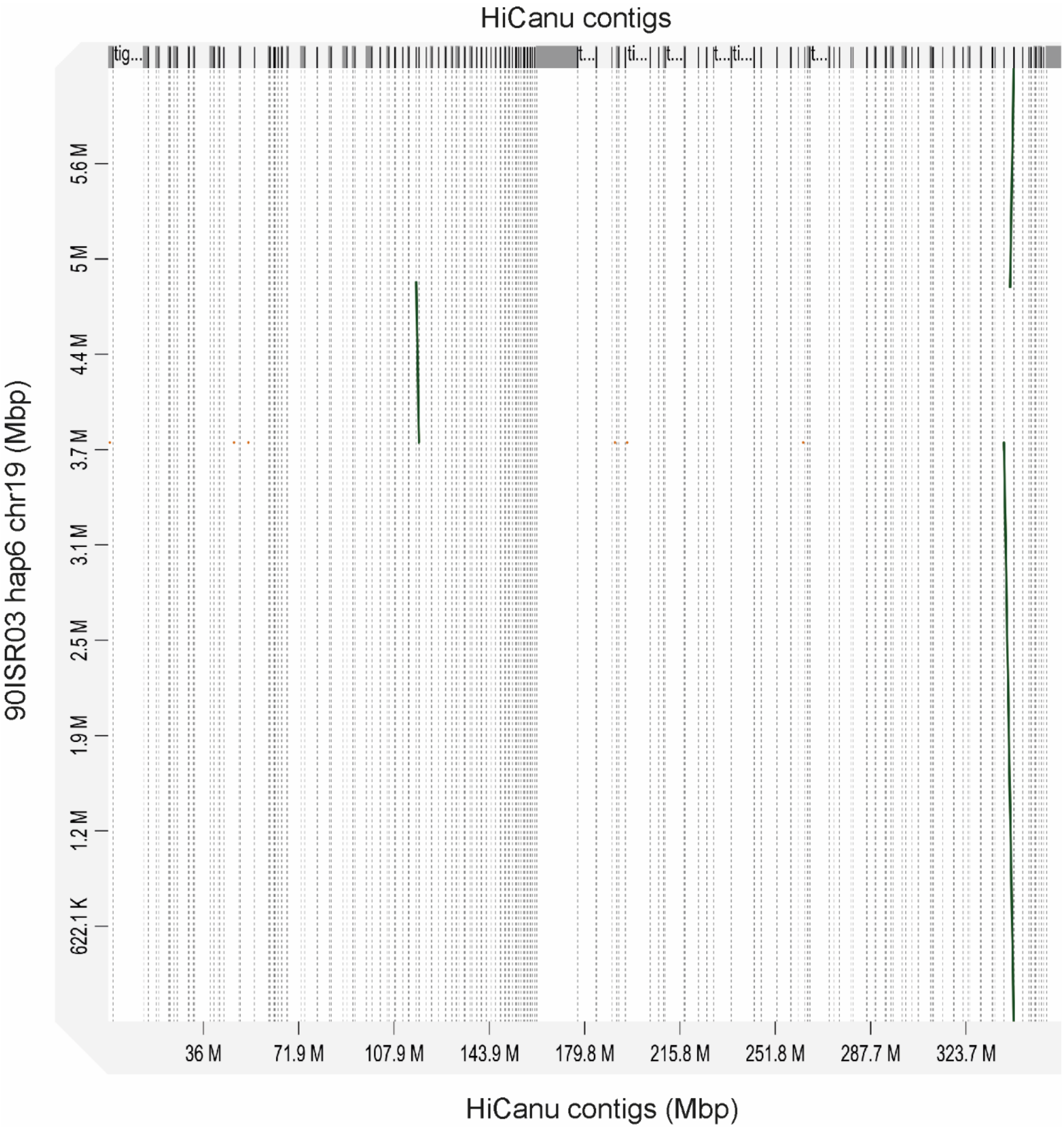


**Figure S13**. D-GENIES plot of pairwise alignment of 90ISR03 hap6 chr19 to the 90ISR03 HiCanu assembly.

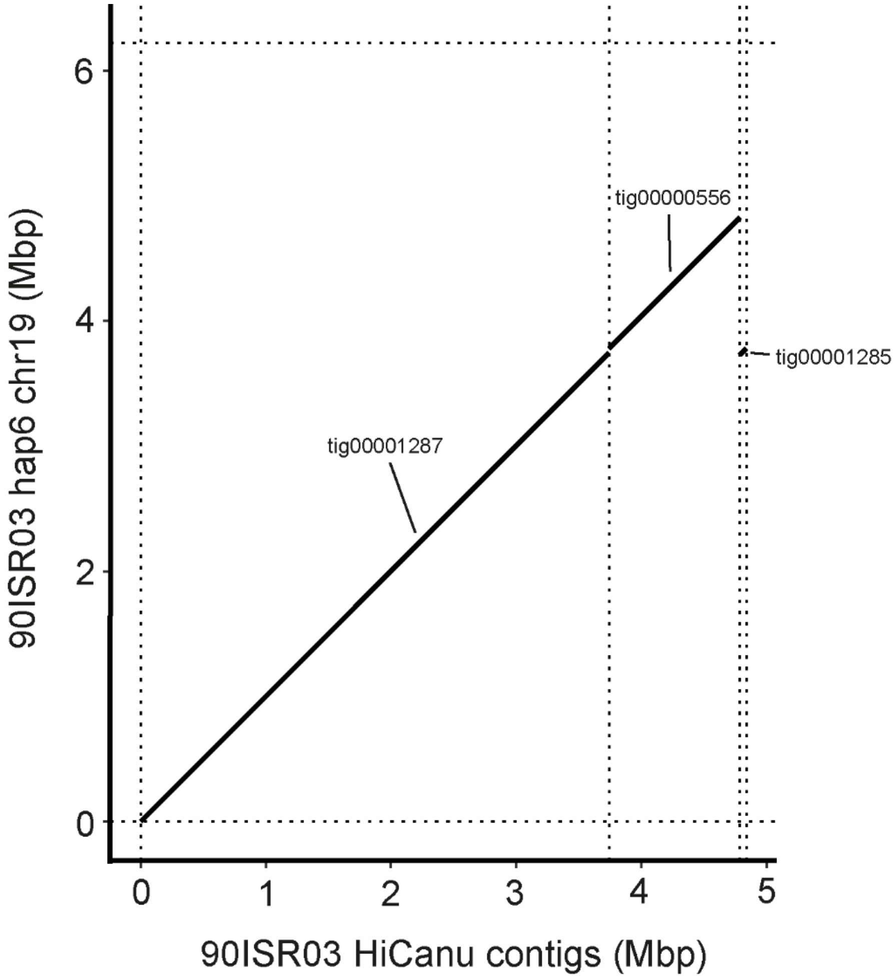


**Figure S14**. D-GENIES plot of pairwise alignment of 90ISR03 HiCanu assembly contigs tig00001287, tig00000556 and tig00001285 to chr19.
